# Mechanisms driving epigenetic and transcriptional responses of microglia in a neurodegenerative lysosomal storage disorder model

**DOI:** 10.1101/2024.11.12.623296

**Authors:** Christopher D. Balak, Johannes C.M. Schlachetzki, Addison J. Lana, Elizabeth West, Christine Hong, Jordan DuGal, Yi Zhou, Benjamin Li, Payam Saisan, Nathanael J. Spann, Vishal Sarsani, Martina P. Pasillas, Sydney O’Brien, Philip Gordts, Beth Stevens, Fredrik Kamme, Christopher K. Glass

## Abstract

Lysosomal dysfunction is causally linked to neurodegeneration in many lysosomal storage disorders (LSDs) and is associated with various age-related neurodegenerative diseases^1,2^, but there is limited understanding of the mechanisms by which altered lysosomal function leads to changes in gene expression that drive pathogenic cellular phenotypes. To investigate this question, we performed systematic imaging, transcriptomic, and epigenetic studies of major brain cell types in *Sgsh* null (KO) mice, a preclinical mouse model for Sanfilippo syndrome (Mucopolysaccharidosis Type IIIA, MPS-IIIA)^3,4^. MPS-IIIA is a neurodegenerative LSD caused by homozygous loss-of-function (LoF) mutations in *SGSH* which results in severe early-onset developmental, behavioral, and neurocognitive impairment^5–15^. Electron microscopy, immunohistochemistry, and single-nucleus RNA-sequencing analysis revealed microglia as the cell type exhibiting the most dramatic phenotypic alterations in *Sgsh* KO mice. Further temporal analysis of microglia gene expression showed dysregulation of genes associated with lysosomal function and immune signaling pathways beginning early in the course of the disease. *Sgsh* deficiency similarly resulted in increases in open chromatin and histone acetylation at thousands of putative microglia-specific enhancers associated with upregulated genes but had much less impact on the epigenetic landscapes of neurons or oligodendrocytes. We provide evidence for dominant and context-dependent roles of members of the MITF/TFE family as major drivers of microglia-specific epigenetic and transcriptional changes resulting from lysosomal stress that are dependent on collaborative interactions with PU.1/ETS and C/EBP transcription factors. Lastly, we show that features of the transcriptomic and epigenetic alterations observed in murine *Sgsh* deficiency are also observed in microglia derived from mouse models of age-related neurodegeneration and in human Alzheimer’s disease patients, revealing common and disease-specific transcriptional mechanisms associated with disease-associated microglia phenotypes.

## Introduction

Pathological variants in genes that result in LSDs are a leading cause of childhood neurodegeneration and dementia^16,17^, indicating that defects in the ability of lysosomes to execute their degradative functions are sufficient to drive pathological processes that result in neuronal dysfunction and death. Beyond the degradation of proteins, lipids, carbohydrates and nucleic acids, lysosomes act as metabolic and inflammatory signaling hubs involved in functions that vary in a cell specific manner, including autophagy, energy homeostasis, inflammation and phagocytosis^18–20^. Accordingly, autophagosomal and endolysosomal dysfunction has been implicated in age-related neurodegenerative diseases (reviewed in ^2,21–24^) such as Alzheimer’s disease (AD)^25–28^, Parkinson’s disease (PD)^29,30^, frontal temporal dementia (FTD) and amyotrophic lateral sclerosis (ALS)^31–36^. Similarly, genetic variants causal of neurodegenerative diseases are often associated with the expression or function of genes involved in endolysosomal functions, including *BIN1* in AD^37–39^, *LRRK2* in PD^40–42^, and *GRN* in FTD^43,44^, and *C9ORF72* in ALS^45–47^.

Beyond genetic causes, lysosomal dysfunction can result from local environmental perturbations, including uptake of protein aggregates such as β amyloid or pathogenic forms of tau that are resistant to degradation or alter lysosomal integrity^48^.

In LSDs, all cells bear the pathogenic variant, yet there is limited knowledge of the consequences of the loss of a specific, non-redundant lysosomal degradative capacity across different brain cell types. This deficiency is expected to have some degree of cell autonomous effects in neurons, but neuronal dysfunction and death may also result from effects in other cell types, such as microglia, which rely on robust lysosomal degradative capacity to degrade autophagic and phagocytic cargos. For example, recent studies in mouse models of the LSDs Gaucher disease (GD) and neuronal ceroid lipofuscinosis (NCL) suggest a critical role for microglial lysosomal dysfunction in neuroinflammation and neurotoxicity^49,50^. In GD mice, biallelic LoF mutations in *Gba1* lead to lysosomal accumulation of glycosphingolipids which were found to exert potent inflammatory and immunogenic activities in microglia, while in NCL mice, null progranulin levels via *Grn* KO shifted microglia into a disease-specific state that caused endolysosomal dysfunction and neurodegeneration. However, transcriptional mechanisms by which LoF of a lysosomal enzyme result in altered gene expression in different brain cell types have not been established for any LSD. Elucidation of these mechanisms may provide insights into the consequences of lysosome dysfunction not only in LSDs but also in age-related neurodegenerative diseases.

While most age-related neurodegenerative diseases are complex diseases, LSDs, which are caused by pathogenic variants in lysosomal genes, offer a window of opportunity to study specific lysosomal dysfunction across different brain cell types. One LSD having a profound impact on the brain is MPS-IIIA, caused by LoF mutations in *SGSH* leading to diminished activity of the lysosomal enzyme sulfamidase^14,15^. Children with MPS-IIIA develop progressive behavioral, motor, and cognitive symptoms including neurocognitive impairment and dementia, ultimately succumbing to the disease by late adolescence^5–13^. The inactivation of sulfamidase leads to the lysosomal accumulation of its glycosaminoglycan substrate heparan sulfate (HS), a major component of the brain extracellular matrix (ECM) which microglia actively phagocytose and remodel^4,51–54^. In humans and mice, secondary storage products that become elevated include GM2/GM3 gangliosides and cholesterol^3,4,55–58^, with further characterization in mouse models demonstrating amyloid, tau, and alpha synuclein accumulation in the brain reminiscent of age-related neurodegenerative disorders^59^. Despite significant efforts, no curative or significant disease-modifying therapies are currently available and recent enzyme replacement strategies have not led to measurable clinical benefits^60^. Thus, understanding the underlying disease pathogenesis is needed to develop alternative interventional strategies.

## Results

### Microglia are the most severely affected cell type in the brain of *Sgsh* KO mice

To investigate the impact of *Sgsh* deficiency on gene expression within the brain we performed single-nucleus RNA sequencing (snRNA-seq) on forebrain tissue derived from *Sgsh*^+/+^ (WT), *Sgsh*^+/-^ (Het), and *Sgsh*^-/-^ (KO) mice with late-stage disease (8 months). A total of ∼32,500 high quality nuclei were recovered corresponding to 18 main cell populations after clustering as depicted in the uniform manifold approximation and projection (UMAP) plot in Fig. 1a. Following cell type identification by reference-based label transfer^61^, multiple neuronal cell types could be discriminated which clustered according to neurotransmitter type and anatomical brain regions (Fig. 1a, Extended Data Fig. 1a). A large cluster of oligodendrocytes was identified, whereas nuclei derived from microglia and astrocytes were underrepresented in comparison to their frequency in the brain. After combining smaller neuronal clusters into major inhibitory (GABAergic, *Gad2*^+^), excitatory (glutamatergic, *Slc17a7*^+^), and undetermined classes (Extended Data Fig. 1b), we performed differential gene expression analysis between disease (KO) and unaffected (WT, Het) samples in clusters containing >500 nuclei using NEBULA^62^. Using an absolute fold change (FC) of 1.5 (p.adj<0.05) cutoff for identifying differentially expressed genes (DEGs), microglia were the most significantly affected cell type exhibiting over 530 dysregulated genes, 339 of which were upregulated (Fig. 1b). In contrast, neurons and oligodendrocytes exhibited ∼3-5 fold fewer DEGs, most of which being downregulated in their respective classes. Upregulated genes in microglia included *Apoe*, *Cd9*, *Ctsd*, and *Mitf*; genes commonly found to be upregulated in human neurodegenerative diseases and their corresponding mouse models, whereas downregulated genes included those typically associated with a homeostatic phenotype (*P2ry12*, *Sall1*) (Fig. 1c). Oligodendrocyte DEGs included upregulated *Cadm1* and *Igfr1* and downregulated *Tubb4a*, *Cnp*, and *Plp1*, genes similarly found to be dysregulated in the neurodegenerative contexts of AD and/or mouse models of demyelination^63,64^. These results are consistent with recent findings in both age-related (*e.g.*, PD) and lysosomal-associated (Mucopolysaccharidosis Type II, MPS-II) neurodegenerative disorders wherein microglia exhibit significantly higher gene dysregulation than neurons, oligodendrocytes, or astrocytes^65,66^.

**Figure 1.**
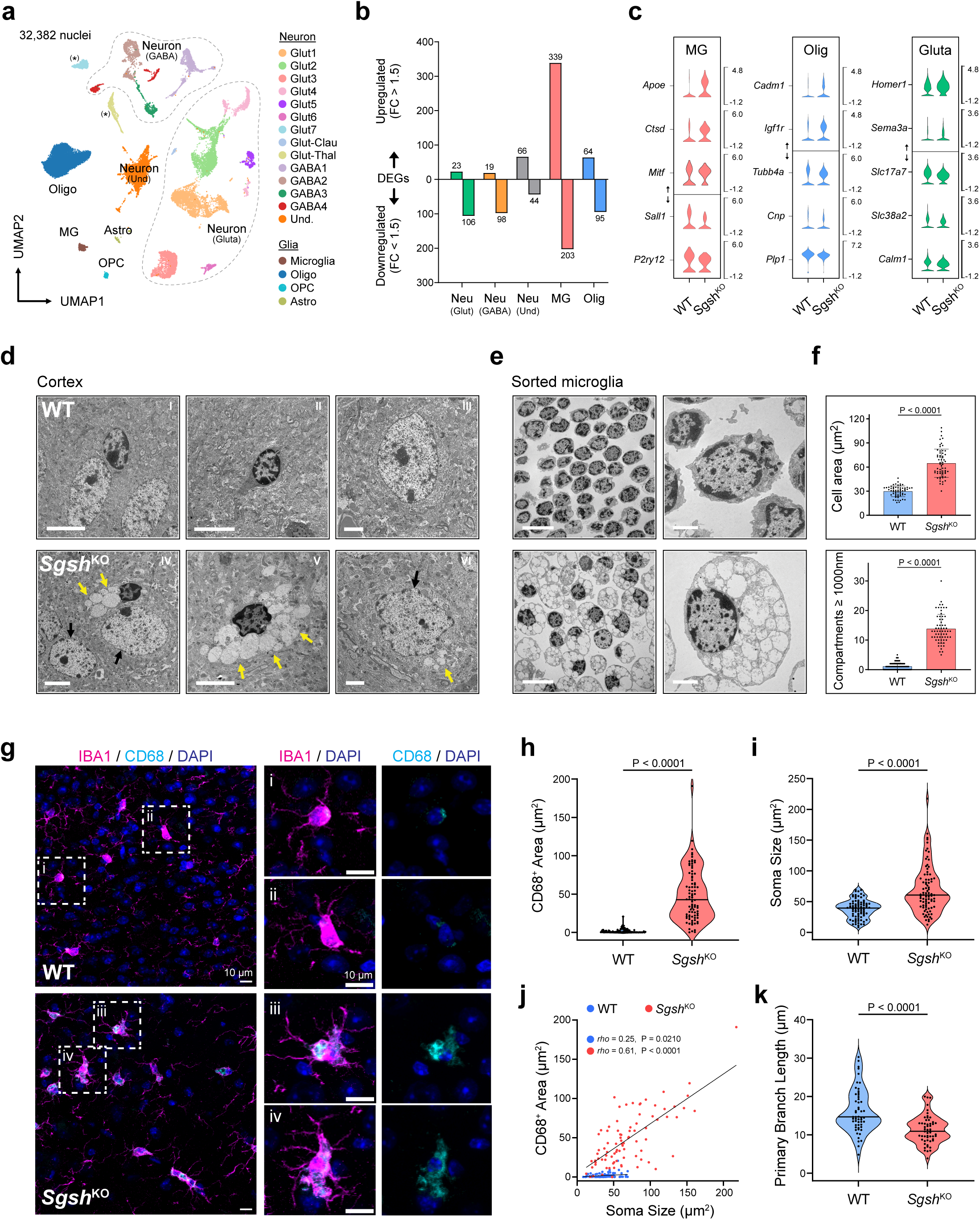
Microglia are the most severely affected cell type in brains of *Sgsh* KO mice. **(a)** UMAP plot of all high-quality captured nuclei (*n=32,382*) from WT, Het, and *Sgsh* KO mice (*n=2* biological replicates/genotype) at 8 months of age. Dashed circles denote major neuronal subclasses grouped by major neurotransmitter usage (glutamatergic (*Slc17a7*^+^) and GABAergic (*Gad2*^+^)). Asterisks denote *Slc17a7*-expressing neurons not clustering with main group. **(b)** Total DEGs (FC≥1.5, FDR≤0.05, NEBULA) in major brain cell types/clusters containing ≥500 nuclei. Smaller neuronal clusters from Figure 1a were combined based on major neurotransmitter type. The neuron-like cluster not displaying clear neurotransmitter identity (Neuron “Und.”) was analysed as its own group for differential expression analysis. **(c)** Violin plots of representative DEGs in microglia (MG), oligodendrocytes (Olig), and glutamatergic neurons (Gluta). All genes reach statistical significance (FDR≤0.05, NEBULA). **(d)** Representative EM images of cerebral cortex sections from WT and *Sgsh* KO mice at 8 months of age (*n=4* biological replicates/genotype). Scale bars: left, 5 μm; middle, 5 μm; right; 2 μm. Yellow arrows denoted expanded, likely lysosomal, cytoplasmic compartments. **(e)** Representative EM images of sorted microglia from WT and *Sgsh* KO brains at 6 months of age (*n=3* biological replicates/genotype). Scale bars: left, 10 μm; right, 2 μm. **(f)** Quantification of sorted microglia EM images. <u>Top</u>: Bar plot illustrating significant increases in cell areas (μm^2^) in *Sgsh* KO microglia. <u>Bottom</u>: Bar plot illustrating significant increases in cytoplasmic compartments >1000 nm in *Sgsh* KO microglia. **(g)** Representative immunofluorescence (IF) imaging in WT and *Sgsh* KO cortex brain sections stained for pan-microglia marker IBA1 (cyan), CD68 (green), and DAPI (blue) at 6 months of age. All scale bars, 10 μm. **(h-k)** Quantification of IF imaging. (**h,i**) Violin plots showing increased Cd68^+^ staining and soma size (as quantified by Iba1 staining) in microglia (*n=2* biological replicates/genotype). Each dot represents an individual cell. Solid lines represent median values, p values determined by unpaired two-tailed t-tests. (**j**) Scatter plot showing correlation of soma size and CD68^+^ staining. Each dot represents an individual cell. P values determined by Spearman’s rank correlation. (**k**) Violin plot of microglia primary branch length quantified by Iba1^+^ staining (*n=2* biological replicates/genotype). Each dot represents the average primary branch length per microglia. Solid lines represent median values, p values determined by unpaired two-tailed t-test.

To assess the consequences of *Sgsh* KO on ultrastructural features of the brain, we analysed the frontal cortex of WT and *Sgsh* KO mice by transmission electron microscopy (EM). These studies revealed large vacuolar structures (>1000 nm diameter) that were most strikingly associated with small, electron-dense nuclei exclusively in KO mice, consistent with lysosomal enlargement in microglia (Fig. 1d subpanel IV-V, yellow arrows, Extended Data Fig. 1c). In contrast, nuclei exhibiting morphological features consistent with neurons, astrocytes and oligodendrocytes were either associated with small vacuolar structures (Fig. 1d subpanel VI, yellow arrow) or had no associated vacuoles (black arrows). To establish whether the cells containing large vacuolar structures associated with small electron-dense nuclei indeed corresponded to microglia, *ex vivo* microglia from WT and *Sgsh* KO mice were sorted and similarly analysed by EM. Sorted microglia exhibited small electron-dense nuclei, with *Sgsh* KO microglia containing large vacuolar structures like those observed *in vivo* (Fig. 1e). Quantitative analysis indicated that the cell area of *Sgsh* KO microglia was approximately twice that of WT microglia (Fig. 1f, upper panel) and uniquely contained numerous cytoplasmic vacuoles >1000 nm in size (Fig. 1f, lower panel). Intriguingly, microglia isolated from *Sgsh* KO mice also exhibited intracytoplasmic inclusions resembling dystrophic neurites^67^ at low frequency, suggesting recent phagocytosis and/or reduced digestion of damaged neuronal substructure (Extended Data Fig. 1d). Further, sorted microglia from *Sgsh* KO mice exhibited increased signal intensities for CD45 and Lysotracker Red by flow cytometry compared to WT or Het mice, consistent with a reactive phenotype and increased acidic organelle content (Extended Data Fig. 1e,f). Supporting these ultrastructural and flow cytometry results, immunofluorescence microscopy examining the pan-microglial marker IBA1 and the reactive microglia-associated phagolysosomal marker CD68 demonstrated markedly increased CD68 staining in *Sgsh* KO microglia (Fig. 1g, h). Quantitative analysis further indicated significant increases in microglia soma size (Fig. 1i) which positively correlated with CD68-positive area (Fig 1j), consistent with EM analysis of sorted microglia. In addition, we observed a significant decrease in primary branch length of microglia processes consistent with reactive microglia (Fig. 1k). Together, these data demonstrate significant and preferential defects in microglia in the *Sgsh* KO brain at both the transcriptional and morphological level.

### Time-dependent changes in microglia gene expression in *Sgsh* KO mice

Although microglia exhibited the most DEGs in the snRNA-seq experiment that examined all brain cell types (Fig. 1a), the relatively low number of microglia recovered limited the depth of analysis that was possible for this cell type. To gain further insights into the effect of *Sgsh* KO in microglia, we performed single-cell RNA sequencing (scRNA-seq) on sorted CD45^+^ cells from WT and *Sgsh* KO mice. Following quality control filtering, 4,021 cells were clustered yielding 10 populations (Extended Data Fig. 2a,b), ∼85% of which were microglia expressing high levels of marker genes *Tmem119* and *P2ry12*, while the other populations were comprised of other immune populations, including macrophages/monocytes, B cells, T cells, and neutrophils. Microglia were further subclustered into 5 populations which exhibited significant differential representation across WT and KO cells and segregated considerably by genotype (Fig. 2a). Most notably, 94% of WT microglia belonged to clusters 2 and 3 compared to only 1% of KO microglia (Fig. 2b). These clusters were associated with a homeostatic molecular signature as indicated by high expression levels of *Tmem119*, *P2ry12, and Cx3cr1* (Extended Data Fig. 2c). Conversely, 83% of *Sgsh* KO microglia, but <1% of WT microglia, resided in cluster 1 (Fig. 2b). This cluster was associated with a reactive DAM-like molecular signature as indicated by high expression levels of *Apoe*, *Cst7*, *Cd9*, *Ctsb*, and *Cd63* (Extended Data Fig. 2c). Subsequent differential expression analysis between WT and *Sgsh* KO cells in the microglia subcluster identified more than 1000 differentially regulated genes at a FC>1.5 (p.adj<0.05). These results suggested that bulk RNA-seq of sorted microglia would be capable of detecting changes in gene expression resulting from *Sgsh* KO due to the large fraction of cells that are affected.

**Figure 2.**
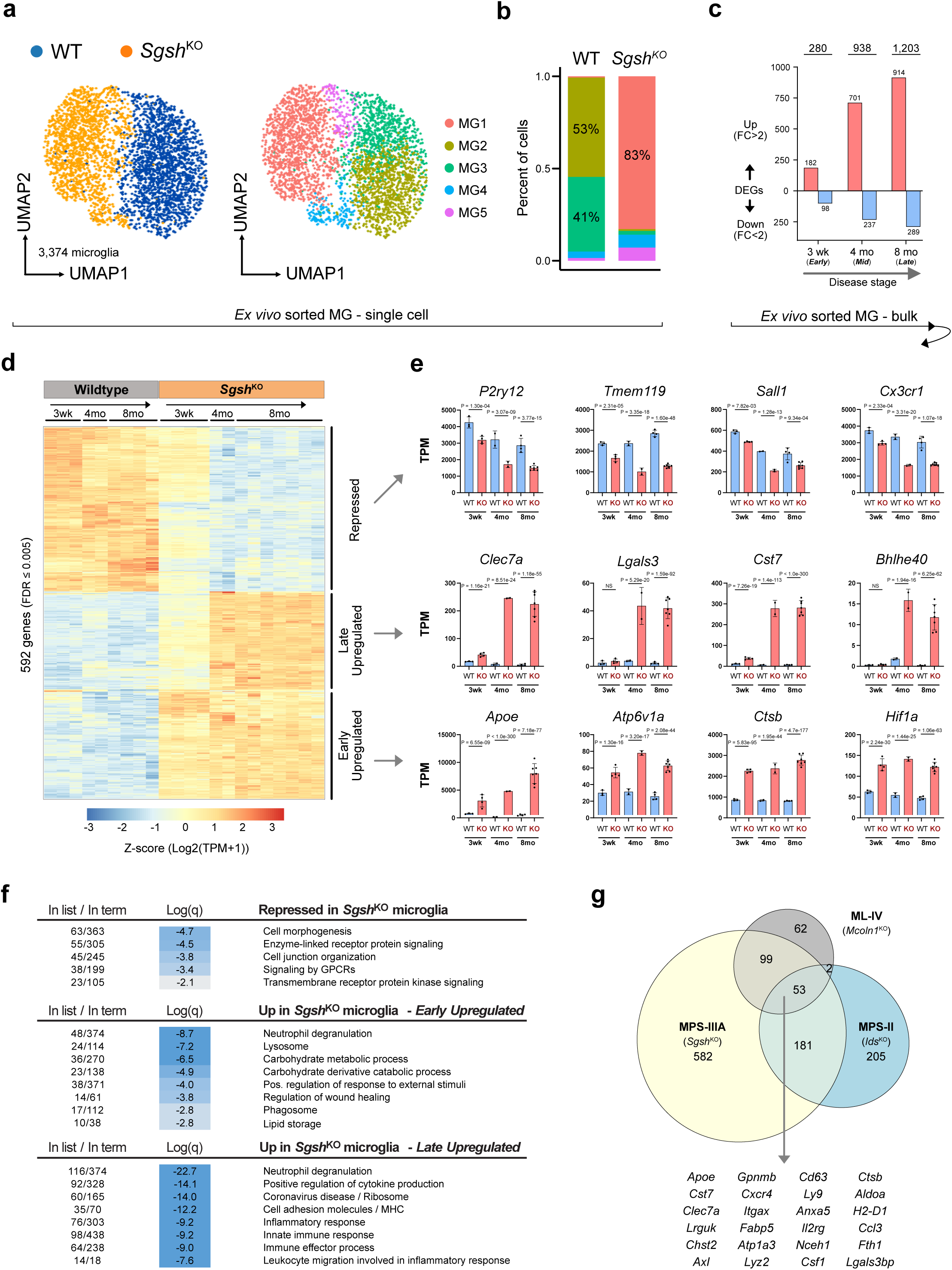
Gene expression signature of *Sgsh* KO microglia. **(a)** UMAP plots annotated for cell genotype (left) and cluster (right) from scRNA-seq in WT and *Sgsh* KO CD45^+^ cells at 6 months of age (*n=2* biological replicates/genotype). **(b)** Bar plot of cluster proportions across genotypes. **(c)** Total DEGs (FC>2, p.adj<0.05, DeSeq2) determined from bulk RNA seq analysis of sorted microglia from WT and *Sgsh* KO mice at the indicated time points (*n=3,4,2,2,4,7* biological replicates for 3wk^WT^, 3wk^KO^, 4mo^WT^, 4mo^KO^, 8mo^WT^, 8mo^KO^, respectively). Red, upregulated genes; blue, downregulated genes. **(d)** Heat map displaying the TPM counts (log_2_(TPM+1)) of the most significant differentially regulated genes across all timepoints (p.adj<0.005, DeSeq2). **(e)** Bar plots of expression levels (TPM) of downregulated, early upregulated, and late upregulated genes in WT and *Sgsh* KO microglia across timepoints. Blue, WT; Red, KO. Data are represented as mean ± standard deviation, p.adj values from DESeq2 analysis (Benjamini– Hochberg method). **(f)** Metascape pathway enrichment analyses of Downregulated (top), Early Upregulated (middle), and Late Upregulated genes (bottom). Repressed: FC<-1.5, p.adj<0.05 (8mo^WT^/8mo^KO^), Early Upregulated: FC>1.5, p.adj<0.05 (3wk^KO^/3wk^WT^ and 8mo^KO^/8mo^WT^), Late Upregulated: FC>1.5, p.adj<0.05 (8mo^KO^/3wk^KO^). **(g)** Venn diagram demonstrating overlap of upregulated genes in *Sgsh* KO microglia with upregulated genes in microglia from LSD mouse models of ML-IV and MPS-II.

We therefore performed RNA sequencing on sorted microglia (CD11b^+^/CD45^low^/CX3CR1^+^ cells, Extended Data Fig. 2d) from WT and *Sgsh* KO mice across disease progression (3 weeks, 4 months, and 8 months of age). These studies revealed a time-dependent increase in DEGs in KO microglia, with a clear molecular phenotype observed even at 3 weeks. Using a FC cutoff of >2 (p.adj<0.05), upregulated genes progressed from 182 genes at 3 weeks to more than 1,200 genes at 8 months of age, while downregulated genes progressed from 98 genes to more than 289 over the same time interval (Fig. 2c, Supplementary Table 1). A heat map illustrating the 592 most significant differentially regulated genes (p.adj<0.005) is illustrated in Fig. 2d. Approximately 40% of these genes exhibited progressive downregulation from 3 weeks to 4 months and remained repressed at 8 months. Downregulated genes included genes typically associated with microglia identity and/or a homeostatic state, including *P2ry12*, *Tmem119*, *Sall1,* and *Cx3cr1* (Fig. 2e, top row). Pathway enrichment analysis of this gene set indicated modest enrichment for terms including cell morphogenesis, enzyme-linked receptor signaling, GPCRs and cell junction organization (Fig. 2f, top).

Two general classes of upregulated genes could be discriminated in *Sgsh* KO microglia. The first, characterized by early upregulation at 3 weeks which remained upregulated at 8 months, included genes such as *Apoe*, *Atp6v1a*, *Ctsb*, and *Hif1a* (Fig. 2e, bottom row). Pathway enrichment analysis of this gene set returned terms including neutrophil degranulation (a gene set enriched for lysosomal enzymes, lysosomal membrane proteins, and vesicular trafficking proteins), lysosome, carbohydrate metabolic processes, and wound healing (Fig. 2f, middle). The second class, characterized by relatively little upregulation at 3 weeks but marked upregulation at 4 months which persisted at 8 months, included genes such as *Clec7a*, *Lgals3*, *Cst7*, *Bhlhe40, Il1b, and Tnf* (Fig. 2e, middle row). Pathway enrichment analysis of this gene set was highly enriched for terms including neutrophil degranulation and inflammatory responses (Fig. 2f, bottom). Notably, there was a progressive increase in the number of upregulated genes with functional annotations linked to inflammatory response and positive regulation of cytokine production from 3 weeks to 8 months of age (Extended Data Fig. 2e).

To place these findings in the context of recently established microglia phenotypes in mouse models of LSDs resulting from the deficiency of iduronate-2-sulfatase (*Ids*, the cause of MPS-II (Hunter syndrome)^68–70^) and loss of function of mucolipin 1 (*Mcoln1*, the cause of Mucolipidosis type IV (ML-IV)^71,72^), we compared the upregulated genes in each model^65,73^ (Fig. 2g). Genes significantly upregulated in *Sgsh* KO microglia at 8 months of age overlapped with more than half of the upregulated genes in microglia in both the MPS-II and ML-IV models, with a core set of 53 genes upregulated in all three models (FC>2, p.adj<0.05). Interestingly, ML-VI and MPS-II showed almost no overlap beyond this core set of genes, indicating both common and distinct programs of dysregulated microglia gene expression resulting from these mutations.

### Altered epigenetic landscapes in *Sgsh* KO microglia suggest transcriptional drivers

To investigate transcriptional mechanisms contributing to changes in gene expression observed in specific brain cell types of *Sgsh* KO mice, we defined open chromatin using the assay for transposase accessible chromatin (ATAC-seq)^74^ and enhancer activity using chromatin immunoprecipitation and sequencing (ChIP-seq) for acetylation of histone H3 lysine 27 (H3K27ac) in multiple brain cell types at 4- and 8-months of age. H3K27ac results from the recruitment of transcriptional coactivators that possess histone acetyltransferase activity, such as p300, and is strongly correlated with enhancer and promoter activity^75^. High quality data sets for ATAC-seq and H3K27ac ChIP-seq assays were obtained from microglia, neurons, and oligodendrocytes (Supplementary Table 2). However, similar to snRNA-seq, astrocytes could not be efficiently recovered and analysed and thus were not investigated further. We then defined reproducible ATAC peaks in microglia, neurons, and oligodendrocytes using the Irreproducible Discovery Rate (IDR) method^76^ and used these peak centers as coordinates for annotation of ChIP-seq data to obtain genome-wide active enhancer and promoter landscapes across cell types. Examples of genomic locations of ATAC-seq and H3K27ac ChIP-seq signals demonstrating specificity of nuclear sorting for microglia, neurons, and oligodendrocytes are provided in Extended Data Fig. 3a.

We compared the extent of H3K27ac associated with ATAC-seq peaks in WT and *Sgsh* KO nuclei for microglia, neurons, and oligodendrocytes at the 4- and 8-month time points to identify genomic loci affected by *Sgsh* KO. Examples of genomic regions exhibiting altered open chromatin and H3K27ac activity in the vicinity of the *Atp6v1a*, *Apoe* and *Hpse* genes are illustrated in Fig. 3a. Using criteria of a >1.5 FC in local acetylation (p.adj<0.05), we identified nearly 6,000 ATAC peaks that exhibited differential H3K27ac in microglia at 4 months of age (Figure 3b). In contrast, only 82 peaks reaching these criteria were identified in oligodendrocytes and none were identified in neurons (Fig. 3b). We interpret the lack of differential H3K27ac in neurons and oligodendrocytes to reflect the much smaller effect of *Sgsh* deficiency on gene expression in these cell types, possibly due to differential dependencies on phagocytic and lysosomal pathways as compared to microglia, and the greater heterogeneity of neurons that would reduce power for observing changes that are limited to specific neuronal subsets.

**Figure 3.**
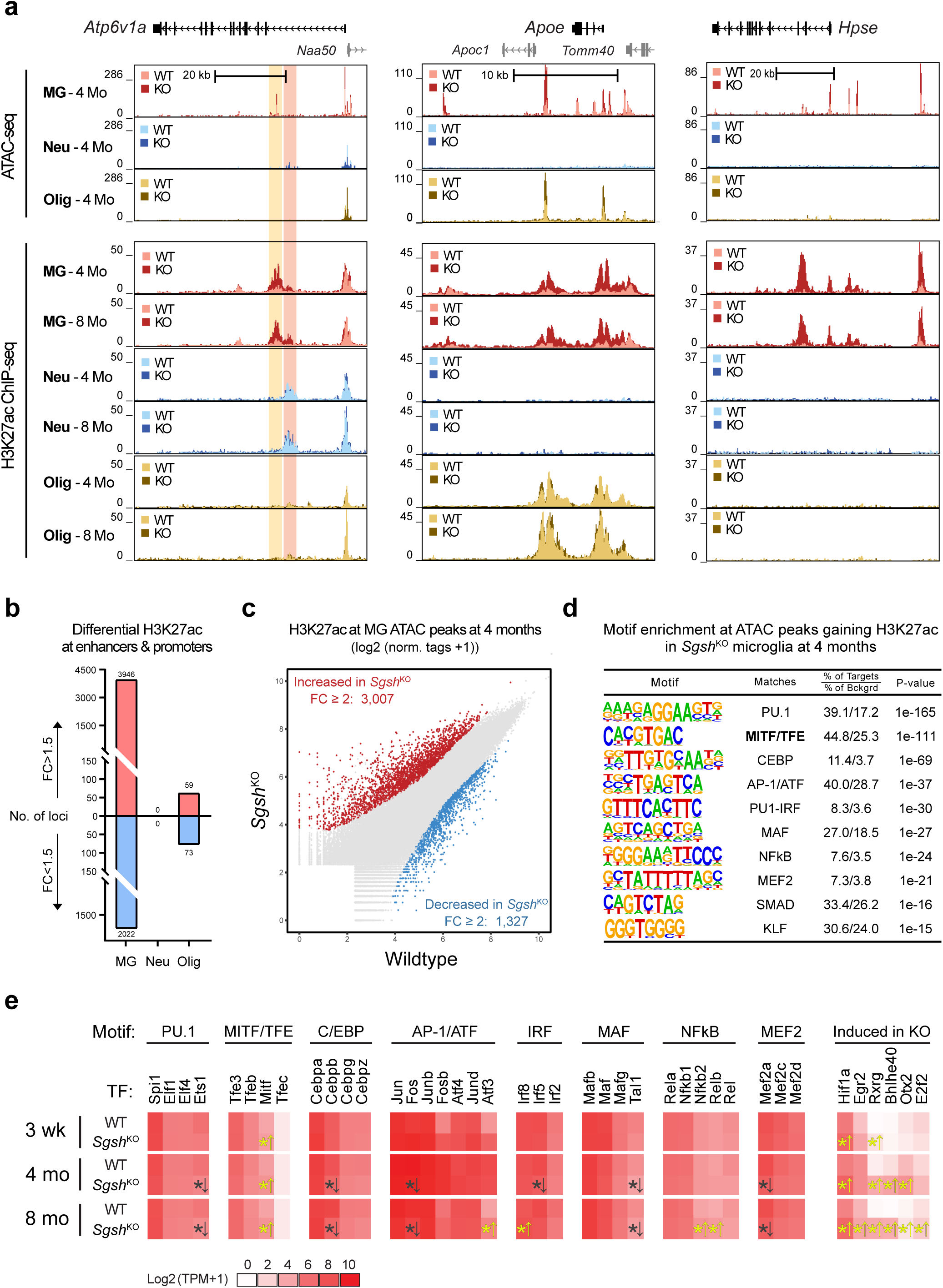
Cell-specific effects of *Sgsh* KO on the epigenetic landscapes of microglia, neurons, and oligodendrocytes. **(a)** Representative ATAC-Seq and H3K27ac ChIP-seq data in UCSC Genome Browser for WT and *Sgsh* KO microglia (MG), neurons (Neu) and oligodendrocytes (Olig) at 4 and 8 months (Mo) of age at the indicated loci (*n=2* biological replicates per genotype and per timepoint in each cell type). **(b)** Number of ATAC-seq peaks exhibiting significant increases or decreases in local H3K27ac in microglia, neurons and oligodendrocytes derived from WT vs *Sgsh* KO mice at 4 months of age. **(c)** Scatter plot of mean H3K27ac (log_2_(norm.tags+1)) at ATAC-seq peaks in WT and *Sgsh* KO microglia isolated at 4 months of age. Red: increased FC>2 (p.adj<0.05) in *Sgsh* KO microglia, blue: decreased FC>2 (p.adj<0.05) in *Sgsh* KO microglia. **(d)** *De novo* enrichment analysis of ATAC peaks exhibiting >2-fold increase in H3K27ac deposition (p.adj<0.05) in *Sgsh* KO microglia at 4 months of age. **(e)** Heat map displaying the TPM counts (log_2_(TPM+1)) of the most highly expressed TFs recognizing motifs identified by *de novo* motif enrichment analysis in (d). Asterisks denote significant changes in expression of ± >35% (p.adj <0.01, DeSeq2) and arrows indicate direction of dysregulation (yellow = upregulated).

A substantial number of genes that were highly and selectively upregulated in *Sgsh* KO microglia were associated with large-scale changes in open chromatin and H3K27ac exclusively in this cell type, as exemplified by the promoter and putative enhancers associated with the *Hpse* gene (Fig. 3a, right). However, we also observed striking specificity in epigenetic responses at genes that are co-expressed across cell types but only activated in *Sgsh* KO microglia. For example, *Atp6v1a* is expressed in all three cell types but is selectively upregulated in microglia in *Sgsh* KO mice. A putative microglia- specific intronic enhancer in this gene (indicated by a vertical orange stripe) shows increased H3K27ac in KO mice at 4 and 8 months of age, but an adjacent putative neuron-specific enhancer (indicated by a vertical light red stripe) exhibited a constant level of H3K27ac (Fig. 3a, left). Additionally, *Apoe* is expressed in both microglia and oligodendrocytes, but in KO mice is only significantly upregulated in microglia. Consistent with this, a shared putative enhancer region associated with *Apoe* in microglia and oligodendrocytes exhibited a constant level of H3K27ac in oligodendrocytes, but strongly increased acetylation in microglia (Fig. 3a, middle).

Approximately 90% of the ATAC peaks exhibiting significantly increased or decreased H3K27ac resided more than 3 kb from promoters, suggesting that the dominant impact of *Sgsh* KO on gene expression resulted from effects on distal enhancers. To gain insights into the relationship between epigenetic changes and gene activation in *Sgsh* KO microglia, we applied a more stringent criteria of a 2-fold increase in H3K27ac at ATAC- seq peaks, yielding 3,007 differential peaks at 4 months of age (red points in Fig. 3c) and 3,348 peaks at 8 months of age (Extended Data Fig. 3b). Approximately 60% of these regions in *Sgsh* KO microglia were confidently present in WT microglia (>16 normalized tags), as exemplified by the loci illustrated in Fig. 3a. This observation suggests that the transcriptional consequences of *Sgsh* KO are in large part due to modifications of the activity states of a pre-existing enhancer landscape, but there is also substantial evidence for selection of *de novo*, context-specific enhancers.

To identify candidate TFs mediating changes in histone acetylation, we performed *de novo* motif enrichment analysis of peaks gaining acetylation in *Sgsh* KO microglia in comparison to a GC-matched random genomic background^77^. This analysis yielded more than a dozen significantly enriched motifs at each time point, of which those that could be most confidently linked to a known TF recognition motif are illustrated in Fig. 3d and Extended Data Fig. 3c. The most significantly enriched motif in these regions in 4- and 8- month-old *Sgsh* KO mice is recognised by the macrophage and microglia LDTF PU.1, consistent with its major role in driving the selection of microglia-specific enhancers through collaboration with additional microglia LDTFs^77,78^. Evidence for such collaboration is provided by significant enrichment of motifs recognised by C/EBP, AP-1, MEF2 and MAF factors. Notably, the second most significantly enriched motif at both the 4 and 8-month time points is nearly identical to the Coordinated Lysosomal Expression and Regulation (CLEAR) element that is preferentially recognised by members of the MITF/TFE subfamily of basic HLH transcriptional regulators, suggesting a coordinated lysosomal response in these cells^79,80^. In addition, an NF-κB motif that is enriched at 4 months became more significant at the 8-month time point (Extended Data Fig. 3c), consistent with the greater upregulation of inflammatory gene expression in 8-month-old *Sgsh* KO microglia. Lastly, a motif recognised by interferon-regulatory factors (IRFs) also became significant at the 8-month time point (Extended Data Fig. 3e). ATAC-seq peaks exhibiting loss of H3K27ac in *Sgsh* KO microglia were found to contain a much more limited set of enriched motifs that were dominated by variations in PU.1 and MEF2 motifs (Extended Data Fig. 3d). This result could be partly due to the smaller number of ATAC peaks exhibiting loss of H3K27ac, limiting statistical power for motif discovery, but may also reflect more complex mechanisms resulting in reduced acetylation that are not readily captured by motif enrichment analyses.

The most highly expressed members of each class of TFs capable of recognizing the motifs identified by motif enrichment analysis are illustrated in Fig. 3e. Many factors did not exhibit significant changes in mRNA expression in *Sgsh* KO microglia, suggesting that consequences of *Sgsh* KO on histone acetylation result from post-transcriptional regulation of TF activity in addition to regulation at the transcriptional level. It is also important to note that *de novo* motif enrichment analysis does not capture all functionally relevant motifs contributing to altered H3K27ac and downstream gene expression and that additional TF families are likely to contribute to these changes. For example, a small number of TFs were upregulated more than 2-fold in *Sgsh* KO microglia at 3-week, 4- and/or 8-month time points for which enriched motifs were not recovered by *de novo* motif enrichment analysis, including *Hif1a, Egr2*, *Rxrg,* and *Bhlhe40* (Fig. 3e), but which nevertheless may be involved in the overall transcriptional response. Consistent with this, interrogation of the ATAC-seq peaks exhibiting gain of H3K27ac in *Sgsh* KO microglia for all known TF motifs present in the JASPAR database^81,82^ recovered factor-specific versions of all the motifs illustrated in Fig. 3d and Extended Data Fig. 3e, but also identified motifs of lower significance that are recognised by several other TFs that are expressed in microglia, including motifs that are recognised by HIF1α, EGR2, RXRγ, and BHLHE40 (Extended Data Fig. 3e).

### Cell specific consequences of *Sgsh* KO on MITF/TFE factors

The observation that the second most enriched motif in ATAC-seq peaks gaining significant H3K27ac in *Sgsh* KO microglia was nearly identical to the CLEAR element recognised by the MITF/TFE family of TFs strongly implicated one or more of these factors in driving the transcriptional response to *Sgsh* KO. As shown in Fig. 3d, *Mitf*, *Tfeb*, and *Tfe3* were expressed above 16 transcripts per million (TPM) in microglia in all conditions evaluated, whereas *Tfec* was below 1 TPM. Further, *Mitf* exhibited increased expression at all time points (39%, 51%, and 79% at 3 weeks, 4, and 8 months, respectively) in *Sgsh* KO microglia. However, transcriptional activities of the MITF/TFE family are largely controlled by post-translational modifications, including phosphorylation, that regulate nuclear entry and protein-protein interactions^83,84^. To investigate the genome wide binding properties of the MITF/TFE family, we developed ChIP-seq assays for each endogenous factor in informative cell lines. We then performed ChIP-seq experiments for each expressed factor in sorted nuclei from mid to late-stage disease (4-8 months) WT and *Sgsh* KO microglia and oligodendrocytes. IDR peak calling yielded high quality peak sets MITF, TFEB, and TFE3 in WT and *Sgsh* KO genotypes for both cell types.

Representative examples of ChIP-seq peaks for MITF, TFEB and TFE3 at promoters and putative enhancer elements associated with *Olig1* (oligodendrocyte-specific), *Apoe* (oligodendrocyte and microglia expressed), and *Hpse* (microglia-specific) are shown in Fig. 4a. At a genome-wide level, we identified thousands of reproducible binding sites for MITF, TFEB and TFE3 in each cell type in WT and *Sgsh* KO mice (Extended Data Fig. 4a, b). *Sgsh* KO resulted in significant increases in the binding of all three factors at most genomic binding sites in microglia and oligodendrocytes, consistent with known MITF/TFE nuclear translocation during conditions of cellular or lysosomal stress^85,86^ (accompanying manuscript, Tejwani *et al*.). However, high magnitude changes in binding (FC>16) were almost exclusively observed in microglia (Extended Data Fig. 4c, d). Within microglia, there was near complete overlap of the binding profiles of MITF and TFEB with the larger profile of TFE3 peaks (Extended Data Fig. 4e). This pattern was similar in oligodendrocytes, although to a slightly less degree than observed in microglia (Extended Data Fig. 4f). This observation is consistent with the proposed mechanism by which MITF/TFE factors function as exclusive hetero- and homodimers to bind their target regions of DNA^87,88^.

**Figure 4.**
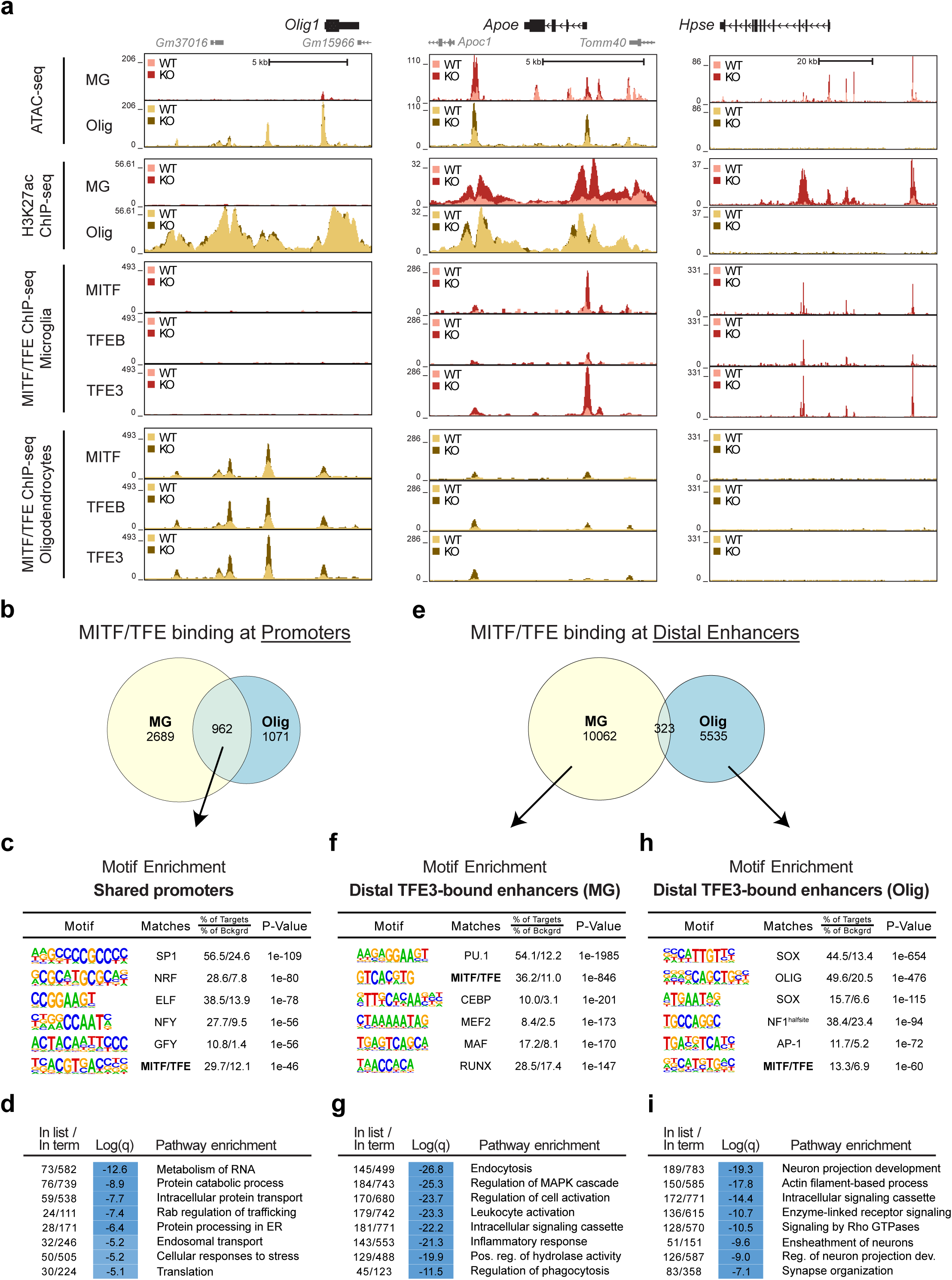
Cell-specific and genotype-dependent binding profiles of MITF, TFEB and TFE3 in WT and *Sgsh* KO microglia and oligodendrocytes. **(a)** Representative ATAC-Seq and ChIP-seq data in UCSC Genome Browser for H3K27ac, MITF, TFEB and TFE3 in WT and *Sgsh* KO microglia (MG) and oligodendrocytes (Olig) at the indicated loci (*n=2-3* biological replicates per genotype in each cell type). **(b)** Overlap of active TFE3 binding sites in *Sgsh* KO microglia and oligodendrocytes at promoters (±500 bp from TSS). Active binding sites were defined as IDR peaks having ≥64 normalized tags for TFE3 and ≥16 normalized merged tags for H3K27ac. **(c)** *De novo* motif enrichment at sites of shared binding of TFE3 at promoters in microglia and oligodendrocytes. **(d)** Metascape pathway enrichment analysis of genes associated with shared binding of TFE3 at promoters in microglia and oligodendrocytes. **(e)** Overlap of active TFE3 binding sites at distal genomic regions (>500 bp up/downstream from TSS) in microglia and oligodendrocytes in *Sgsh* KO microglia. **(f)** *De novo* motif enrichment at microglia-specific active TFE3 binding sites. **(g)** Metascape pathway enrichment analysis of nearest genes to microglia-specific TFE3 binding sites. **(h)** *De novo* motif enrichment at oligodendrocyte-specific TFE3 binding sites. **(i)** Metascape pathway enrichment analysis of nearest genes to oligodendrocyte-specific TFE3 binding sites.

As the binding profile of TFE3 included the large majority of the binding sites of MITF and TFEB in each cell type, we considered its binding at promoters and distal genomic regions to be representative of the binding programs of all three factors. As expected, there was a significant degree of overlap of TFE3 binding at promoters in the two cell types (Fig. 4b). These regions were enriched for motifs recognised by general promoter-biased transcription factors, including SP1, ELF and NFY, in addition to the CLEAR motif recognised by the MITF/TFE factors (Fig. 4c). The associated genes were enriched for annotations linked to basic cellular functions including RNA metabolism, protein degradation and intracellular protein transport (Fig. 4d). In stark contrast, the binding program of TFE3 at putative enhancers in *Sgsh* KO mice was largely cell type specific (Fig. 4e). For example, at the *Olig1* locus, binding of all three factors was exclusively observed at the *Olig1* promoter and putative upstream oligodendrocyte-specific enhancers (Fig. 4a, left). At *Hpse*, binding of MITF, TFEB and TFE3 were highly induced selectively at open regions of chromatin that are exclusively present in microglia (Fig. 4a, right). Consistent with this, motif enrichment analysis of TFE3 binding sites at putative microglia enhancers returned the CLEAR element and motifs recognised by microglia LDTFs (Fig. 4f), with the nearest expressed genes showing enrichment for functional annotations linked to immune cell function (Fig. 4g). Conversely, motif enrichment analysis of oligodendrocyte-specific TFE3 peaks at regions of active chromatin returned the CLEAR element and motifs recognised by oligodendrocyte LDTFs (Fig. 4h). The nearest expressed genes to these binding sites were enriched for functional annotations linked to oligodendrocyte function, including ensheathment of neurons (Fig. 4i). These findings suggest that while members of the MITF/TFE family exert shared functions in microglia and oligodendrocytes, they also exert diverse cell-specific functions that are established through collaborative interactions with microglia or oligodendrocyte LDTFs at distal regulatory elements.

### Association of MITF/TFE factors with homeostatic and stress-responsive transcriptional states

The acquisition of the *in vivo* genome-wide binding locations of the MITF/TFE family members in WT and *Sgsh* KO microglia enabled a direct evaluation of the extent to which these factors were associated with changes in the epigenetic landscape and corresponding motif enrichment patterns. Two general categories of MITF/TFE binding were observed for each factor; a category of basal binding under homeostatic conditions that was unchanged or decreased in the *Sgsh* KO, and a category of low or basal binding that was significantly increased in the *Sgsh* KO. An example of a location at which the basal binding of MITF, TFEB and TFE3 is unchanged is indicated by a vertical yellow stripe in an intronic enhancer-like region of the *Hexb* gene (Fig. 5a). This peak is near a location at which there is a varying degree of increased binding of each factor in the *Sgsh* KO, indicated by the vertical blue stripe. However, the level of H3K27ac for the entire region was equivalent between genotypes and the expression of *Hexb* was not changed in *Sgsh* KO microglia. Conversely, an example at which the binding of MITF and TFE3 were strongly increased in the *Sgsh* KO is indicated by a vertical yellow stripe downstream of *Cst7*. Here, there is a marked increase in H3K27ac and *Cst7* was strongly upregulated in *Sgsh* KO microglia (Fig. 5b). A similar pattern is exhibited at the *Hpse* locus (Fig. 4a), which was also strongly upregulated in the *Sgsh* KO.

**Figure 5.**
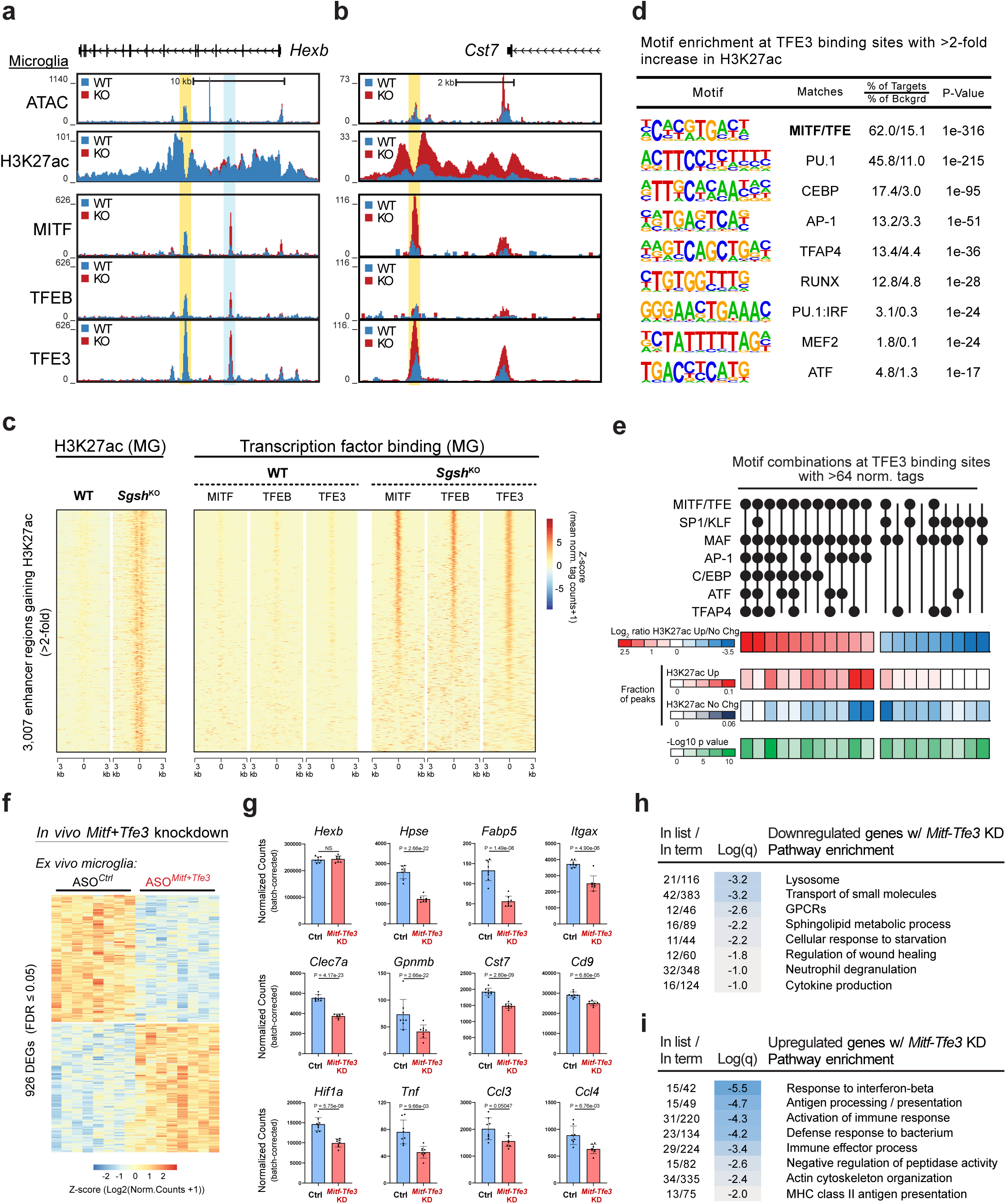
Homeostatic and *Sgsh* KO-dependent binding programs of MITF, TFEB and TFE3 in microglia. **(a)** Representative ATAC-Seq and ChIP-seq data in UCSC Genome Browser for H3K27ac, MITF, TFEB, and TFE3 in WT and *Sgsh* KO microglia in the vicinity of *Hexb*. **(b)** Representative ATAC-Seq and ChIP-seq data in UCSC Genome Browser for H3K27ac, MITF, TFEB and TFE3 in WT and *Sgsh* KO microglia in the vicinity of *Cst7*. **(c)** Tag density distribution at ATAC-seq peaks exhibiting a two-fold increase in H3K27ac in *Sgsh* KO microglia (left), and corresponding MITF, TFEB, and TFE3 tag densities at the same regions across genotypes (right). **(d)** *De novo* motif enrichment analysis at ATAC seq peaks with >64 normalized tags for TFE3 and more than a 2-fold increase in H3K27ac in *Sgsh* KO microglia. **(e)** Motif co-occurrence analysis of TFE3 binding sites associated with gain of H3K27ac or no change of H3K27ac in *Sgsh* KO microglia. **(f)** Heat map of differential gene expression (p.adj<0.05) in *ex vivo* microglia from 3-month-old *Sgsh* KO mice injected intracerebroventricularly with a combination of *Mitf-* and *Tfe3-targeting* ASOs or control ASO for 14 days (*n=8* biological replicates per genotype). **(g)** Representative downregulated genes from (f). All genes displayed are significantly upregulated in *Sgsh* KO microglia compared to WT. *Hexb* is displayed as a control with unchanged expression in between WT and *Sgsh* KO mice. P values are p.adj from DeSeq2. **(h)** Metascape pathway enrichment analysis of downregulated genes. **(i)** Metascape pathway enrichment analysis of upregulated genes.

To investigate the relationship between MITF/TFE binding and changes in H3K27ac at a genome wide level, we plotted the tag densities of H3K27ac, MITF, TFEB and TFE3 ChIP-seq data as a function of the distance from peak centers at the 3,007 regions that exhibited a >2-fold increase in H3K27ac (p.adj<0.05) in *Sgsh* KO microglia (Fig. 5c). This analysis revealed high magnitude increases in the binding of all three factors at most of these regions, analogous to the patterns observed at the *Cst7* and *Hpse* loci, and is consistent with roles of these factors in driving increased histone acetylation at a large fraction of the genomic regions affected by *Sgsh* KO.

These findings indicated that while unaltered binding of MITF/TFE factors between WT and *Sgsh* KO genotypes is generally associated with stable H3K27ac, only a subset of sites exhibiting increased MITF/TFE binding were associated with increased H3K27ac and increased gene expression. To investigate the basis for this observation, we performed *de novo* motif enrichment analysis of TFE3 binding sites (>64 normalized tags) exhibiting a >2-fold increase in H3K27ac (p.adj<0.05) in *Sgsh* KO microglia (similar to the peak marked by the vertical yellow stripe in Fig. 5b and corresponding to the TFE3 peaks in Fig. 5c) in comparison to TFE3 binding sites exhibiting significant (>16 normalized tags) but unchanged (<1.2 FC) H3K27ac (similar to the peak marked by the vertical blue bar in Fig. 5a). At TFE3 binding sites associated with increased H3K27ac, a consensus TFE3 binding site was the most highly enriched motif, present in 62% of target sequences, followed by motifs for PU.1, C/EBP factors, AP-1, and TFAP4/HLH factors (Fig. 5d). In contrast, at TFE3 binding sites associated with unchanged H3K27ac, the consensus TFE3 binding site decreased in significance and was only present in 31% of the peaks (Extended Data Fig. 5). Additional top enriched motifs in this subset included a GC-rich motif recognised by SP-1 and KLF factors (third most enriched) and a motif recognised by the MAF factors.

Based on these results, we performed an iterative analysis of the frequencies of motif co-occurrences of the top motifs identified in each of the two peak sets over a range of motif thresholds. An example of the output of this analysis for MITF/TFE, SP-1/KLF, MAF, AP- 1, C/EBP, ATF and TFAP motifs at an intermediate threshold is illustrated in Fig. 5e. Of the 128 possible combinations of these motifs, 37 exhibited significantly higher representation in one peak set or the other (p<0.05), with the 20 most significantly different combinations (p<0.002) illustrated in Fig. 5e. At the motif thresholds used for this example, all the motif combinations associated with gain of H3K27ac included a MITF/TFE motif, but this was the case for only 3 of the 9 combinations associated with no change in H3K27ac. In addition, motif combinations including AP-1, C/EBP and ATF factors were strongly biased in peaks exhibiting a gain of H3K27ac, whereas motif combinations including SP-1/KLF motifs were strongly biased in peaks exhibiting no change in H3K27ac. This general pattern was robustly observed over a broad range of motif thresholds. PU.1 and MEF2 motifs did not contribute to differential peak bias. Collectively, these findings suggest that both the presence of a consensus MITF/TFE motif and specific combinations of nearby motifs for specific additional factors, including C/EBP, AP-1 and ATF factors, are required for TFE3 binding to result in histone acetylation and increased gene expression.

### Molecular consequences of loss of function of MITF/TFE3 in *Sgsh* deficiency

To further investigate the transcriptional functions of the MITF/TFE factors, we designed anti-sense oligonucleotides (ASOs) targeting mouse *Mitf*, *Tfeb* and *Tfe3* to specifically knockdown their mRNA levels *in vivo*. Single bolus doses of each ASO were delivered intracerebroventricularly into 3-month-old WT mice and microglia were isolated by FACS after 14 days. Initial RNA-seq studies demonstrated significant and specific *in vivo* reductions of mRNA levels of 96%, 87% and 81% for *Mitf*, *Tfeb* and *Tfe3,* respectively (Extended Data Fig. 5b, c). Despite significant DNA binding of MITF, TFEB and TFE3 at thousands of genomic locations in homeostatic/WT microglia, reduced expression of their corresponding single mRNAs in response to ASO treatment was associated with few changes in global gene expression. Given their co-expression and highly overlapping DNA binding patterns, we considered MITF, TFEB and TFE3 might have redundant functions in microglia similar to *in vitro* findings in other cell types thus limiting the consequences of individual knockdowns^86,89^. Additionally, as SDTFs, significant effects might only be observed when these factors are significantly active in the cell, for instance, during conditions of lysosomal stress. We therefore performed *in vivo* knockdown experiments in 3-month-old *Sgsh* KO mice using a combination of ASOs targeting *Mitf* and *Tfe3* (Extended Data Fig. 5d). Due to technical and dosing limitations, it was not possible to inject all three ASOs simultaneously. However, MITF and TFE3 were associated with the most genomic locations in *Sgsh* KO microglia, exhibited the largest changes in DNA binding in *Sgsh* KO microglia, and were the only MITF/TFE family members which significantly translocated to the nucleus in macrophages in the context of another lysosomal disorder (accompanying manuscript, Tejwani *et al*.). The combination of the *Mitf* and *Tfe3* ASOs, which resulted in parallel knockdowns of 80% and 56%, respectively (Extended Data Fig. 5e), resulted in a large number of low magnitude but highly significant changes in mRNA expression in comparison to the control ASO. In total, 460 mRNAs were significantly downregulated and 466 mRNAs upregulated by the combined treatment (p.adj<0.05) (Fig. 5f).

Downregulated genes resulting from *Mitf/Tfe3* knockdown exhibited a high degree of overlap with genes upregulated in *Sgsh* KO microglia, exemplified by *Hpse*, *Fabp5*, *Clec7a*, *Gpnmb*, *Itgax*, *Hif1a,* and *Cst7* (Fig. 5g). Accordingly, these genes were enriched for functional annotations highly similar to those which were enriched in the upregulated gene sets of *Sgsh* KO microglia, including lysosome, neutrophil degranulation, inflammatory response, and regulation of wound healing (Fig. 5h). In contrast, genes exhibiting increased expression in mice treated with the combination of *Mitf* and *Tfe3* ASOs included interferon-inducible genes, such as *Ifi206*, *Isg15*, and MHC class II genes, and were enriched for functional annotations linked to response to interferon β, activation of the immune response and antigen processing (Fig. 5i). Collectively, these findings support redundant roles of MITF/TFE family members in driving a significant component of the transcriptional response to *Sgsh* deficiency.

### Intersection with mouse models of age-related neurodegeneration and human AD

To examine the relevance of these findings to mouse models of age-related neurodegenerative diseases, we intersected genes differentially regulated in *Sgsh* KO microglia with genes differentially regulated in microglia responding to either 1) amyloid-induced brain pathology, or 2) to brain pathology induced by the combination of human P301S tau and APOE4 (Fig. 6a). Responses to amyloid-induced pathology were represented by RNA-seq data comparing CLEC7A^high^ microglia (referred to as Microglia- neurodegeneration; MGnD^90^, a phenotype highly overlapping disease-associated microglia (DAM)^91^) to CLEC7A^low^ microglia (homeostatic microglia; Hom) isolated from APP/PS1 mice at 9 months of age, at which time significant amyloid pathology is present. The amyloid pathology-responsive microglia (AR-MG) phenotype is partially dependent on TREM2 signaling and has been linked to neuroprotective functions of microglia^91–93^. We defined 577 AR-MG genes from this comparison using a FC>2 (p.adj<0.05). In contrast, mice that express the combination of human P301S Tau and APOE4 (TE4 mice) develop severe neurodegeneration, including marked neuronal death and volume loss in the hippocampus^94^. Neurodegeneration in this context is strongly attenuated by treating mice with a CSF1 receptor inhibitor, indicating that microglia play essential roles in driving tissue pathology in this model^95^. Using snRNA-seq, a previously unidentified cluster of microglia was identified in TE4 mice, but not in mice selectively expressing APOE4 (E4 mice), which do not develop neurodegeneration^96^. These cells were referred to as tau/APOE4-Responsive Microglia, or TERM. In comparison to homeostatic microglia, TERM exhibited 733 genes upregulated >2-fold (p.adj<0.05) and only partially overlapped those that are upregulated in AR-MG (CLEC7A^high^) microglia (Fig. 6a).

**Figure 6.**
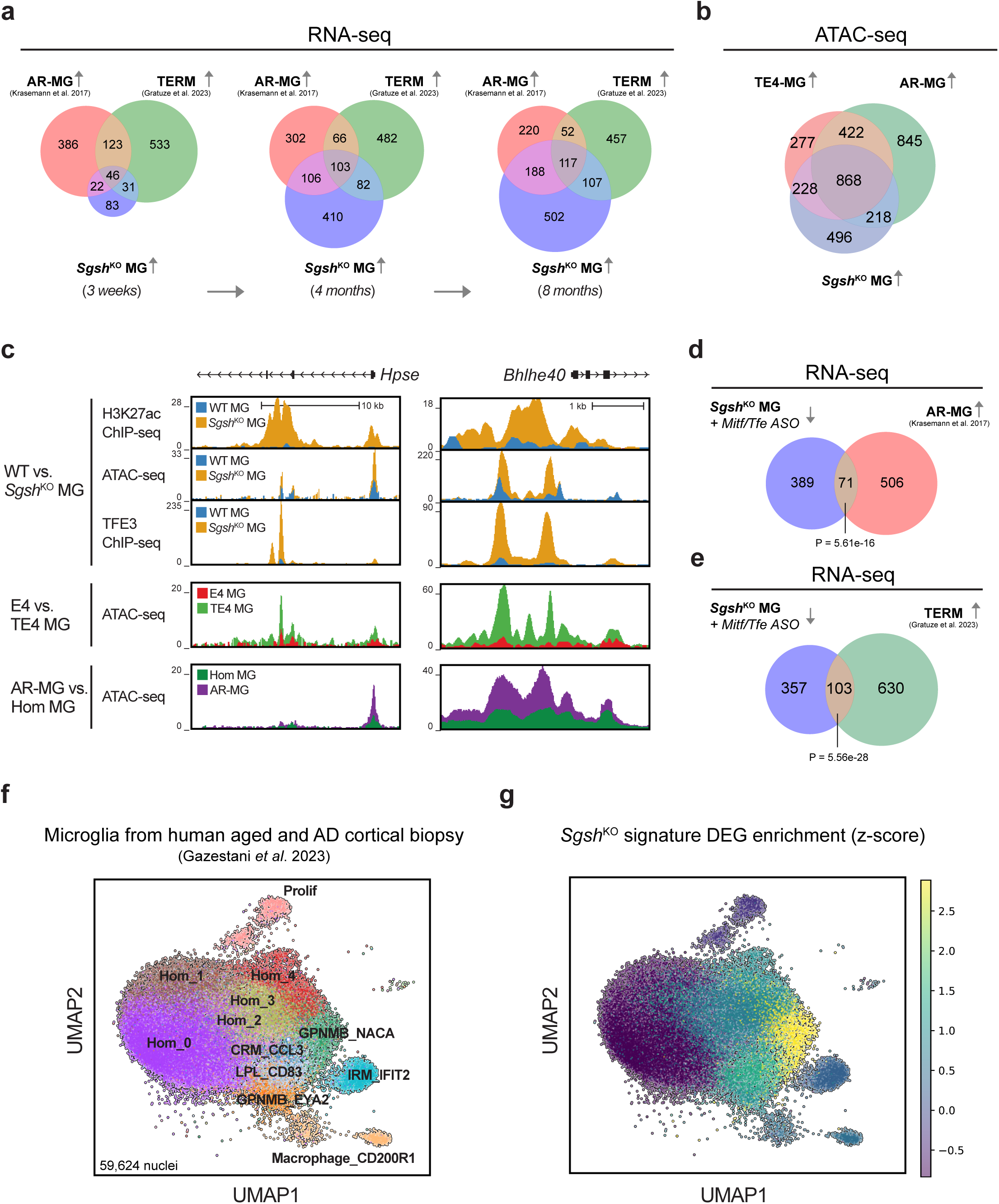
Relationships to models of age-related neurodegeneration and human microglia in AD. **(a)** Intersection of genes upregulated in *Sgsh* KO microglia at 3 weeks (left), 4 months (middle), and 8 months of age (right) (purple) with CLEC7A^high^ amyloid-pathology responsive microglia (AR-MG) from APP/PS1 mice (red) and Tau/apoE-responsive microglia (TERM) from TE4 mice (green). **(b)** Intersection of genomic locations of induced open chromatin (ATAC peaks) in *Sgsh* KO microglia with induced open chromatin in AR-MG and TE4-responsive microglia. **(c)** UCSC Genome Browser tracks of representative ATAC-Seq and ChIP-seq data at the *Hpse* and *Bhlhe40* loci in *Sgsh* KO vs WT microglia, AR-MG vs homeostatic microglia (Hom), and TE4 vs E4 microglia. **(d)** Intersection of genes downregulated in *Sgsh* KO microglia treated with a combination of *Mitf* and *Tfe3* ASOs (p.adj<0.05) and mRNAs upregulated in AR-MG (FC>2, p.adj<0.05). P values: Fisher’s exact test. **(e)** Intersection of genes downregulated in *Sgsh* KO microglia treated with a combination of *Mitf* and *Tfe3* ASOs (p.adj<0.05) and mRNAs upregulated in TERM (FC>2, p.adj<0.05). P values: Fisher’s exact test. **(f)** UMAP plot of the cell clusters of brain myeloid cell clusters obtained from snRNA-seq of human cortical biopsies from individuals with varying degrees of amyloid and tau pathology as published in Gazestani *et al*. 2023. Each dot represents a single cell. **(g)** UMAP plot of the composite averaged z-score from human orthologs of a representative *Sgsh* KO signature gene set across the cells in the UMAP shown in **f**.

Of the 182 genes upregulated in *Sgsh* KO microglia at 3 weeks of age, 68 genes overlapped with the upregulated AR-MG gene set and 77 with the upregulated TERM microglia gene set, of which 46 were in common (Fig. 6a, left). This overlap expanded at 8 months of age, with 305 of the 914 genes upregulated in *Sgsh* KO microglia intersecting with the upregulated AR-MG gene set (corresponding to 53% of AR-MG genes), and 224 with the TERM set (corresponding to 31% of TERM genes) (Fig. 6a, right). Notably, only 117 of the overlapping genes were common to all three microglia phenotypes. Thus, there was a distinct segregation of genes upregulated in the *Sgsh* KO microglia phenotype with those associated with a protection-associated, amyloid-pathology responsive signature and a neurodegeneration-associated TE4-responsive signature.

As described in the accompanying manuscript (Schlachetzki/Hasselmann *et al*.), we used ATAC-seq to define the AR-MG and TE4-responsive open chromatin environments in microglia at the same time points used to assess differential gene expression in the APP/PS1 and TE4 models. Although ATAC-seq is less sensitive than H3K27ac in defining changes in regulatory elements, it was possible to define 1,494 regions of open chromatin in AR-MG and 917 regions of open chromatin in TE4 microglia that exhibited >2-fold more tags (p.adj<0.05) than corresponding control microglia (Hom in the case of AR-MG and E4 in the case of TE4 microglia). Performing a similar analysis for *Sgsh* KO microglia, we identified 944 regions of open chromatin that exhibited >2-fold more tags (p.adj<0.05) than in WT microglia. Alignment of these regions indicated both a high degree of overlap in the induced open chromatin environment in all three model systems, with 868 shared peaks, but also substantial model-specific peaks (Fig. 6b). Examples of common *Sgsh* KO-, amyloid- and TE4-responsive regions of open chromatin aligned with H3K27ac data in *Sgsh* KO and WT mice are illustrated at the *Hpse* and *Bhlhe40* loci in Fig. 6c. Notably, at subsets of these sites there are high magnitude increases in TFE3 binding in *Sgsh* KO microglia. In addition, there is reduced expression of a subset of these genes in *Sgsh* KO mice treated with the *Mitf*+*Tfe3* ASO combination (Fig. 5g, 6d, e). These results are consistent with common mechanisms driving selected aspects of the altered open chromatin and gene expression changes observed in *Sgsh* KO mice, in responses to amyloid pathology, and responses to pathogenic tau and APOE4 due to lysosome dysfunction.

In order to investigate whether the gene expression changes observed in our *Sgsh* KO microglia are reflected in human microglia in AD, we conducted an analysis using snRNA-seq data from two recent large-scale studies of human microglia from the prefrontal cortex. These studies included individuals across the AD continuum, encompassing both individuals at various stages of disease and those without AD. The first, Gazestani *et al*.^97^, consisted of snRNA-seq data from the fresh biopsies of 52 individuals, and the second, Green *et al*.^98^, included snRNA-seq data from post-mortem samples from 437 individuals. To assess the overlap between the *Sgsh* KO gene signature and human patient-derived microglia, we generated a gene signature score derived from ∼350 highly significant and upregulated genes in *Sgsh* KO microglia. This gene list was then assigned aggregated enrichment z scores in the human nuclei data sets. This analysis revealed that the *Sgsh* KO DEG signature was most highly enriched in the GPNMB_NACA subpopulation followed by the LPL_CD83 and GPNMB_EYA2 subpopulations in the fresh biopsy cohort, together comprising the three reactive and AD-enriched clusters defined in the dataset (Fig. 6f, g). In the post-mortem dataset, the *Sgsh* KO signature was found to be highly enriched in Mic.10 (reactive redox), Mic.13/Mic.12 (Lipid-associated disease subpopulations), Mic.15 (inflammatory), and to a lesser extent in Mic.14 (interferon response) (Extended Data Fig. 7a, b). These findings provide evidence that the microglia response to altered lysosomal function via *Sgsh* KO significantly overlaps with the human microglial response to AD pathology in human brains, both in fresh biopsy and post-mortem tissue.

## Discussion

In contrast to common and sporadic forms of age-related neurodegenerative diseases, in which altered transcriptional states are concentrated in subsets of cells that are localized to focal areas of pathology, germline monogenic *Sgsh* KO has the potential to exert cell autonomous consequences in all cell types in which it is expressed. Unexpectedly, ultrastructural, immunohistochemical, transcriptional and epigenetic analyses indicate that microglia are by far the most affected cell type in the brain parenchyma of *Sgsh* KO mice. Indeed, we observed abundant gene expression changes in microglia, contrasting the relatively few changes in neurons, oligodendrocytes or other brain cell types. These findings provide evidence that microglia play dominant roles in mediating uptake and clearance of extracellular SGSH substrates including HS and raise questions as to mechanisms by which homozygous LoF mutations result in neurodegeneration. The relatively modest changes in gene expression and morphology of other brain cell types coupled with the dramatic lysosomal expansion and transcriptional changes in the microglia lysosomal gene expression program are consistent with a compensatory protective function of microglia in mitigating pathogenic effects of HS accumulation in other brain cell types early in the course of disease. We speculate that neuropathology ensues when the capacity of microglia and their lysosomes to perform this function eventually becomes overwhelmed, resulting in a combination of increased cell autonomous effects of HS accumulation in neurons and other brain cell types and neurotoxic effects of the altered program of gene expression in microglia. The transition in the functional annotations of differentially regulated genes to an increasingly inflammatory signature between the 3-week and 8-month time points is consistent with this possibility. Based on this interpretation, early administration of cell replacement therapies in which *SGSH*-mutant microglia are replaced with WT microglia or emerging brain-penetrant enzyme replacement therapies have potential for therapeutic benefit in disorders affecting microglia or multiple CNS cell types^65,99^.

The primary objective of these studies was to investigate transcriptional mechanisms through which LoF of a non-redundant, degradative lysosomal enzyme results in altered gene expression within cell types of the brain through an analysis of altered epigenetic landscapes. Quantitative analysis of open chromatin and H3K27ac identified substantial changes in microglia, but significantly fewer in neurons or oligodendrocytes, consistent with gene expression data. The altered epigenetic profile identified in microglia in this context may represent a general signature of a microglia lysosomal stress response, such as the one identified in the accompanying study investigating *Grn* deficiency (Tejwani *et al*.). It will therefore be of interest to expand analysis of epigenetic changes in other LSDs and in contexts in which lysosomes are damaged, such as by uptake of cholesterol or monosodium urate crystals.

The finding that a motif recognised by members of the MITF/TFE family of TFs was among the most significantly enriched motifs in promoters and putative enhancers exhibiting increased H3K27ac activity in *Sgsh* KO microglia was of particular interest because of the established roles of these factors in the regulation of lysosomal function and biogenesis, autophagy, phagocytosis and immune responses^79,80,84–86,100,101^.

Genome wide location analyses of MITF, TFEB and TFE3 in oligodendrocytes and microglia demonstrated a high degree of overlap of binding of these factors within each cell type, but highly divergent binding patterns across cell types, particularly at putative distal microglia- or oligodendrocyte-specific enhancers. At these locations, motif enrichment analysis indicated very similar MITF/TFE motifs in both cell types, but different co-enriched motifs that represented DNA recognition elements for microglia-specific or oligodendrocyte-specific LDTFs, respectively. These findings are consistent with a model in which the genomic binding sites for MITF/TFE factors are determined in each cell type by collaborative interactions with combinations of microglia or oligodendrocyte-specific LDTFs and additional more broadly expressed SDTFs, which collectively enable cell-specific signal-dependent effects on gene expression. Consequently, MITF/TFE factors are likely to play significantly different biological roles in microglia and oligodendrocytes beyond their shared roles in regulation of lysosomal biology.

In both microglia and oligodendrocytes, binding sites for MITF/TFE factors could be broadly partitioned into genomic locations at which their binding was increased in the *Sgsh* KO or genomic locations at which there was significant binding in WT cells that was unchanged or decreased in the knockout. Unexpectedly, in the case of oligodendrocytes, sites of increased binding of MITF/TFE factors were rarely associated with increased H3K27ac and nearby gene expression. In addition, only a subset of genomic regions exhibiting increased binding of MITF/TFE factors in microglia was associated with increased H3K27ac and nearby gene expression. These observations suggest an important additional layer of regulation beyond nuclear translocation that is required for gene activation. Motif co-occurrence analysis suggests that this may be determined at least in part by the specific combinations of co-binding factors and suggests unexpected complexities in gene activation by MITF/TFE factors that may be linked to both protective and pathological functions.

Regarding the association of lysosomal disfunction with age-related neurodegenerative diseases, it was of particular interest to observe the overlap in genes upregulated in microglia in *Sgsh* deficiency with genes that are upregulated in in microglia responding to amyloid- or tau/APOE4-induced pathology, a relationship that was also observed at the level of altered chromatin accessibility. Further, at these locations, there was a high degree of overlap with binding sites for MITF/TFE factors, providing evidence that mechanisms underlying the transcriptional response to lysosomal stress resulting from *Sgsh* deficiency may also contribute to the altered epigenetic landscape and program of gene expression of amyloid pathology-responsive MG and TERM (Schlachetzki and Hasselmann *et al*.). A major question for future investigation is whether this response serves neuroprotective or pathogenic roles. To the extent that activation of the MITF/TFE family of transcription factors enhances the ability of lysosomes to execute their normal functions and compensate for environmental perturbation-associated neuropathology, the functions of these factors are likely to be protective. However, the collateral activation of genes associated with inflammation may be pathogenic. An important objective will be to determine whether it is possible to influence the context and gene-specific activities of MITF/TFE factors for therapeutic purposes.

## METHODS

### Mice

All animal procedures were approved by the University of California San Diego Animal Care and Use Committee in accordance with the University of California San Diego research guidelines for the care and use of laboratory animals. The following mouse strains were used in this study: C57BL/6J (The Jackson Laboratory, Stock No. 00064) and germline *Sgsh* KO^3^ (Dr. Jeffery Esko laboratory and Dr. Philip Gordts laboratory, University of California, San Diego). Mice were housed at the University of California San Diego animal facility on a 12-hour/12-hour light/dark cycle with free access to normal chow food and water.

### Nuclei isolation for snRNA-seq

Forebrain was dissected from six-month-old WT, Het, and KO mice and olfactory bulbs removed. Dissected tissue was homogenized in lysis buffer (10% Triton X-100, 0.1 M DTT, 40 U/μL RNasin (Promega, N2115), protease inhibitor cocktail (Sigma-Aldrich, P8340), and 2% bovine serum albumin (BSA) (Sigma Aldrich, B6917) in PBS) using a Dounce homogenizer for 10 strokes. Samples were incubated for 5 minutes on ice and then centrifuged at 500xg for 5 minutes at 4°C in a tabletop centrifuge. Pellets were resuspended in sorting buffer (0.5 M EDTA, 40 U/μL RNasin, 2% BSA in PBS), transferred to FACS tubes, filtered through a 70 μm mesh strainer, and centrifuged at 800xg for 5 minutes at 4°C in a swinging bucket centrifuge. DAPI (1:1,000; BioLegend 422801) was used to label nuclei. Samples were strained through a 70 μm mesh strainer, washed once more, resuspended in sorting buffer, and kept on ice until nuclei sorting. Samples were sorted for DAPI^+^ singlets on a Beckman Coulter MoFlo Astrios EQ.

### Single nucleus RNA-seq and analysis

Sorted purified nuclei were centrifuged at 800xg for 5 minutes at 4°C and then resuspended in PBS supplemented with 400 mg/mL BSA. Appropriate volumes of each sample were loaded onto a Chromium Controller (10x Genomics) to couple to beads targeting 20,000 cells per sample per the manufacturer’s Single Cell 3’ HT v3.1 (HT) protocol. Following capture, single nuclei sequencing libraries were prepared following the 10X Genomics HT protocol. Samples were sequenced on an Illumina NovaSeq X platform targeting 50,000 reads per cell.

Raw FASTQ files for each sample were processed through CellRanger v7.2.0 (10x Genomics) with standard parameters aligning to the mm10-2020-A reference transcriptome. Downstream analysis was performed with the single cell analysis suite ‘scRNAsequest’^102^. To remove ambient RNA contamination, CellBender^103^ was used with the command scRMambient using the ‘expected_cells’ values equivalent to the captured cell values from 10x and the ‘droplets_included’ values ∼25-50% higher than the 10x captured cell values to ensure capturing empty droplets. CellBender-corrected h5 files were then ran through the scRNAsequest pipeline with the command scAnalyzer; keeping genes expressed in at least 20 nuclei, containing 100-5,500 genes, a maximum 18,000 UMIs, and a doublet score of less than 0.02. Normalization, dimensionality reduction, and integration were carried out by Seurat^104–106^ and Harmony (v1.2.0)^107^. Cluster/cell type-specific differential gene expression analyses were carried out using NEBULA (v1.5.3)^62^, specifically the NEBULA-HL algorithm. Data visualization of data and figure generation was performed in CellxGene^108^.

### Electron microscopy and analysis

Samples were processed and prepared by the UCSD CMM Electron Microscopy Facility. *<u>For EM on cortex sections</u>*, mice were transcardially perfused with Ringer solution followed by 10-15 mL of Fixative Solution (2% paraformaldehyde / 2.5% glutaraldehyde in 0.15 M sodium cacodylate buffer). Brains were quickly removed, dissected into hemispheres, and post-fixed in Fixative Solution at room temperature for 2-4 hours then overnight at 4°C. They were then incubated in 1% OsO4 in 0.1 M cacodylate buffer for 1 hour on ice and then stained with 2% uranyl acetate for 1 hour on ice. The samples were dehydrated in a graded series of ethanol washes (50%-100%) once, followed by a wash with 100% ethanol and 2 washes with acetone for 15 minutes each, and embedded with Durcupan. 70 nm sections were cut on a Leica UCT ultra-microtome and collected on 300 mesh copper grids. Sections were stained with 2% uranyl acetate for 5 minutes, and Sato lead stain for 1 minute. Samples were viewed using a JEOL 1400-plus TEM (JEOL, Peabody, MA). Transmission electron microscopy images were taken using a Gatan OneView 4kx4k camera (Gatan, Pleasanton, CA). Quantification was performed with the Fiji-ImageJ v1.54f software. For vacuole quantification, cytoplasmic / perinuclear compartments greater than 1000 nm in diameter at their widest cross section were counted in 8-21 microglia or non-microglial cells in each biological replicate for each genotype (WT^MG^=35, WT^NonMG^=43, MG^KO^=27, KO^NonMG^=67 cells). Statistical analyses were formed using one-way ANOVA and post-hoc Tukey multiple comparisons in Prism 10 (GraphPad).

*<u>For EM on sorted ex-vivo microglia</u>*, cells were pelleted and chemically fixed with freshly prepared fixative (2.5% glutaraldehyde, in 0.1 M cacodylate buffer pH 7.4). They were then post-fixed in 1% OsO4 in 0.1 M cacodylate buffer for 1 hour on ice and then stained with 2% uranyl acetate for 1 hour on ice. The samples were dehydrated in a graded series of ethanol washes (50%-100%) once, followed by a wash with 100% ethanol and 2 washes with acetone for 15 minutes each, and embedded with Durcupan. 70 nm sections were cut on a Leica UCT ultra-microtome and collected on 300 mesh copper grids. Sections were stained with 2% uranyl acetate for 5 minutes, and Sato lead stain for 1 minute. Samples were viewed using a JEOL 1400-plus TEM (JEOL, Peabody, MA). Transmission electron microscopy images were taken using a Gatan OneView 4kx4k camera (Gatan, Pleasanton, CA). Quantification was performed with Fiji-ImageJ v1.54f software. For microglia cell areas, 20 cells across 2-4 FOVs were traced and quantified in each replicate for each genotype (WT=60, KO=60 cells). For vacuole quantification, cytoplasmic compartments greater than 1000 nm in diameter at their widest cross section were counted in 20-40 cells in each replicate for each genotype (WT=100, KO=63 cells). Statistical analyses were performed using unpaired two-tailed t-tests in Prism 10 (GraphPad).

### Immunofluorescence and analysis

WT and *Sgsh* KO mice were perfused and fixed with 4% paraformaldehyde in PBS. Samples were sectioned (Leica CM3050 S) at 16 μm and collected on Superfrost Plus slides. Sections were fixed with 4% PFA for 10 min, permeabilized in PBS-T (PBS with 0.3% Triton X-100), and blocked with 5% donkey serum (Sigma, D9663) for 2 hours at room temperature. Slides were incubated with primary antibodies 1:1000 rabbit anti-Iba1 (FujiFilm, 019-19741) and 1:300 rat anti-Cd68 (Bio-Rad, MCA1975) at 4°C overnight, then incubated with 1:500 secondary antibodies (donkey anti-rabbit 555, Invitrogen A32794; donkey anti-rat 650, Invitrogen SA5-10029) for 2 hours at room temperature, counter-stained with DAPI, and mounted with Prolong Gold antifade reagent. Images were acquired on a Leica SP8 Confocal microscope with a 63x objective and processed with ImageJ. 63x z-stack FOVs across cerebral cortices of two mice per genotype were analysed. Raw images were max projected, binarized, and thresholded with Otsu’s Method. Cd68^+^ staining areas within Iba1^+^ microglia were quantified. 85 individual cells were measured over 10-16 FOVs per group. ‘Fill Holes’ command was used to fill in cell somas before measuring microglia soma sizes. Microglia branching was quantified using NeuronJ^109^ measuring 54 individual cells over 6-11 FOVs per group. Statistical analyses were performed using unpaired two-tailed t-tests in Prism 10 (GraphPad).

### Isolation of live microglia

Forebrain from mice were homogenized and microglia isolated as previously described^110,111^. Briefly, forebrains were dissected and olfactory bulbs removed. Tissue was homogenized by gentle mechanical dissociation, filtered, washed, microglia enriched with a Percoll gradient, and washed again. All steps were maintained at 4°C with the exception of the Percoll spin. Cells were then incubated in staining buffer on ice with anti- CD16/32 blocking antibody (BioLegend 101319) for 15 minutes, then with anti-mouse anti-CD11b-APC (BioLegend 101212), anti-CD45-Alexa488 (BioLegend 103122), and anti-CX3CR1-PE (BioLegend 149006) for 20 minutes. Cells were then washed and filtered through a 40 μm cell strainer. Sorting was performed on a Sony MA900 or Beckman Coulter MoFlo Astrios EQ cell sorter. Microglia were defined as events that were DAPI negative, singlets, and CD11b^+^/CD45^low^/CX3CR1^+^. Isolated microglia were then processed according to protocols for RNA-seq, ATAC-seq, Lysotracker or EM.

### Single cell RNA-seq and analysis

CD45^+^ sorted cells from wildtype and *Sgsh* KO mice were spun down at 400xg for 5 minutes, supernatant carefully aspirated, and resuspended to 60 μL PBS supplemented with 400 μg/mL BSA. Appropriate volumes of each sample were loaded onto a Chromium Controller (10x Genomics) to target 10,000 cells per sample. Following capture, single cell sequencing libraries were generated using the Chromium Single Cell 3’ Library Reagent Kit (v3 Chemistry, 10X Genomics) following manufacturer’s instructions. Samples were sequenced on an Illumina NovaSeq platform targeting 50,000 reads per cell.

Raw FASTQ files for each sample were processed through CellRanger v7.2.0 with standard parameters aligning to the mm10-2020-A reference transcriptome. Lower than expected cell capture rates led to mean reads per cell increasing 2 to 3-fold per sample. Resulting filtered matrices were analysed in R v4.2.3 using Seurat v4.4.0^104^. QC metrics were added to the Seurat object using the package scCustomize (v2.1.2). Genes expressed in less than 20 droplets were discarded, and droplets meeting the following criteria were kept for downstream analysis: containing >400 and <9500 genes, UMIs >1500, mitochondrial genes <10%, and a complexity (log_10_(nFeature) / log_10_(nCount)) >0.78 and <0.92. The scDblFinder package was used to identify and remove putative doublets^112^. All samples were processed uniformly through one 10x Chromium Chip and library preparation and sequenced together on one Illumina flowcell. No significant batch effects were observed thus integration was not performed. Normalization and dimensionality reduction were carried out according to the standard Seurat v4 pipeline. Major cell types were determined by clustering and known marker gene analysis. The microglia cluster was isolated and subclustered and the package dittoSeq (v1.8.1) was used for comparing cluster distributions. Differential gene expression analysis between genotypes in the microglia cluster was carried out using NEBULA (v1.5.3)^62^, specifically the NEBULA-HL algorithm.

### Sorted microglia RNA sequencing

Total RNA was isolated from 100-300K sorted microglia with RNeasy Plus micro (Qiagen, 74034) or Direct-zol RNA MicroPrep (Zymo Research, R2052) kits and stored at -80°C until RNA-seq library preparation. RNA-seq libraries were prepared as previously described^77,110^. Briefly, polyA mRNA was enriched by two successive rounds of incubation with Oligo d(T) Magnetic Beads (NEB, S1419S) and then fragmented/eluted by incubation at 94°C for 9 min. cDNA was then synthesized with Superscript III (first-strand synthesis; Life Technologies kit 18080-044) and then DNA Polymerase I (second-strand synthesis; Enzymatics P7050L) with dUTP. Resulting cDNA libraries were prepared for sequencing by blunting/end repair, A-tailing, and barcoded adapter ligation (NEXTflex, Bioo Scientific/PerkinElmer) as previously described in Heinz et al^77^. The resulting RNA-seq libraries were digested by incubation with Uracil DNA Glycosylase for 30 min at 37°C. Libraries were PCR amplified for 11-14 cycles, size selected for 200-500 bp fragments by gel excision, quantified using a Qubit dsDNA HS Assay Kit (Thermo Fisher Scientific) and single-end sequenced on a HiSeq 4000 or NextSeq 500 (Illumina, San Diego, CA) according to the manufacturer’s instructions.

### Bulk RNA-, ATAC-, and ChIP-seq preprocessing and mapping

RNA-seq FASTQ files were mapped using STAR (v2.5.3a)^113^ using the mm10 genome and default parameters. For ASO RNA-seq experiments, FASTQ files were pseudomapped using Kallisto (v0.44.0)^114^ with --single -l 200 -s 30 --rf-stranded --single-overhang -b 50 parameters. ATAC- and ChIP-seq data were mapped with Bowtie2 (v2.3.5.1)^115^, with ATAC-seq reads trimmed to 30bp prior to mapping. After mapping, HOMER (v4.11)^77^ was used to convert reads aligning to a single genomic location into “tag directories” for subsequent analysis using the command makeTagDirectory. HOMER was also used to generate sequencing-depth normalized bedGraph/bigWig files for visualization in the UCSC Genome Browser (http://genome.ucsc.edu). A summary of sequencing QC metrics can be found in Supplementary Tables 3 and 4.

### RNA-seq analysis

Raw gene expression counts from tag directories were quantified by HOMER’s analyzeRepeats command with parameters “-condenseGenes -count exons -noadj”, and TPM counts with parameters ‘‘-count exons -tpm’’. TPM counts were matched to raw counts using accession numbers. Small RNAs (<200 bp) and genes with TPM<4 in all conditions were removed from analysis. Differential gene expression was calculated with the HOMER command ‘getDiffExpression.pl’ utilizing DESeq2^116^ (v1.36.0) and DEGs were accessed at FC>2 and p.adj<0.05 where indicated. For ASO RNA-seq experiments, abundance files from Kallisto pseudoalignment were imported with tximport^117^ (v1.24.0) and processed with DESeq2 (v1.36.0) controlling for batch (design = ∼batch + treatment) and keeping genes where the sum of normalized counts in each treatment group was >50 counts. Resulting filtered matrices were further analysed in R v4.2.3. Heat maps were generated using the package pheatmap (v.1.0.12). Pathway enrichment analysis was performed with Metascape using GO Biological Processes, Reactome gene sets, and KEGG pathways (v3.5)^118^. A summary of RNA-seq sequencing QC metrics can be found in Supplementary Table 3.

### LysoTracker assay on live microglia and analysis

Live sorted microglia were pelleted at 400xg for 5 minutes at 4°C following sorting. Supernatant was removed and the pellets resuspended in 1 mL of RPMI media without Phenol Red (Gibco, 11835-030) prewarmed to 37°C. 100 μL of the microglia in media was removed to serve as the negative control. LysoTracker Red (Invitrogen, DND-99) was added to samples at 1:500 to a final concentration of 2 nM and incubated at 37°C for 30 minutes. Following LysoTracker incubation, samples were spun down, media removed, and cells were resuspended in fresh warm RPMI media without LysoTracker. Samples and non-LysoTracker negative controls were ran through a Sony MA900 measuring fluorescence on the PE-Texas Red channel. Quantification of LysoTracker signal was performed with the FlowJo software (v10.4.1). Subsequent one-way ANOVA and post-hoc Tukey multiple comparisons statistical analyses were performed in Prism 10 (GraphPad).

### Isolation of brain nuclei for ATAC / ChIP-sequencing

Brain nuclei were isolated as previously described^37,119^, with initial homogenization performed with either 1% formaldehyde in PBS for 10 minutes (histone ChIP-seq assays), 0.1% formaldehyde in PBS for 10 minutes (oligodendrocyte and neuron ATAC-seq), or 2 mM DSG (Proteochem) for 30 minutes followed by 1% formaldehyde in PBS for 10 minutes (transcription factor ChIP-seq). Nuclei were stained overnight at 4°C with PU.1- PE (Cell Signaling 81886S), OLIG2-AF488 (Abcam 225099), or NEUN-AF488 (Millipore MAB 377X). Nuclei were washed with 4 mL FACs buffer, passed through a 40 μm strainer, and stained with 0.5 μg/mL DAPI (BioLegend 422801). Nuclei for each cell type were sorted with a Beckman Coulter MoFlo Astrios EQ or BD Influx Cell Sorter and pelleted at 1600xg for 5 minutes at 4°C in FACS buffer. Nuclei pellets were snap frozen and stored at -80°C prior to library preparation.

### ATAC-seq

ATAC-seq libraries were prepared as previously described^37,74,120,121^ with approximately 50,000 sorted microglia. Cells were lysed in 150 µl lysis buffer (10 mM Tris-HCl pH 7.5, 10 mM NaCl, 3 mM MgCl2 and 0.1% IGEPAL CA-630 in water). Resulting nuclei were centrifuged at 500xg for 10 min. For oligodendrocytes and neurons, approximately 200,000 lightly crosslinked sorted nuclei were used. Pelleted nuclei were resuspended in 50 μL transposase reaction mix (47.5 μL ATAC lysis buffer^121^ and 2.5 μL Tagment DNA TDE1 enzyme I (Illumina, 20034197)) and incubated at 37°C for 30 min. DNA was purified with Zymo ChIP DNA concentrator columns (Zymo Research, D5205), eluted with 11 µL of elution buffer, and amplified using NEBNext High-Fidelity 2x PCR Master Mix (New England BioLabs, M0541) with the Nextera primer Ad1 (1.25 µM) and a unique barcoding primer Ad2 (1.25 µM) for 8-12 cycles. Resulting libraries were size selected by gel excision to 155–250 bp, purified and single-end sequenced using a HiSeq 4000 (Illumina) according to the manufacturer’s instructions.

### ChIP-seq

Chromatin immunoprecipitation sequencing was performed as previously described^77,122^. *For H3K27ac ChIP-seq*, ∼500,000 fixed nuclei were quickly thawed on ice and resuspended in ice-cold LB3 buffer (10 mM Tris-HCl pH 7.5, 100 mM NaCl, 1 mM EDTA, 0.5 mM EGTA, 0.1% Na-Deoxycholate, 0.5% N-lauroylsarcosine) and 1x protease inhibitor cocktail. Nuclei / chromatin was sheared by sonication using the PIXUL Multi-Sample Sonicator (Active Motif) in low bind 96 well plates (Corning, #7007) following the manufacturer’s instructions. Samples were sonicated with the following settings: Pulse (N): 50, PRF [kHz]: 1.00, Process time: 60 min, Burst Rate [Hz]: 20.00. Samples were pulse spun to 600xg at 4°C in the 96-well plate, transferred to low-bind tubes, and chromatin was recovered by retaining the supernatant following a spin at 21,000xg at 4°C for 10 minutes. The supernatant was then diluted 1.1-fold with ice-cold 10% Triton X-100. Two percent of the lysate was kept as ChIP input control. 10 μL of Dynabead Protein A and 10 μL of Protein G were added per sample, in addition to 2 μg of a specific antibody for H3K27ac (Active Motif 39685). The samples were rotated overnight at 4°C and were washed as follows the next day: 3x with Wash Buffer I (20 mM Tris-HCl pH 7.5, 150 mM NaCl, 2 mM EDTA, 0.1% SDS, 1% Triton X-100) + 1x protease inhibitor cocktail, 3x with Wash Buffer III (10 mM Tris-HCl pH 7.5, 250 mM LiCl, 1% Triton X-100, 1 mM EDTA, 0.7% Sodium Deoxycholate) + 1x protease inhibitor cocktail, 2x with TET (0.2% Tween- 20/TE) + 1/3x protease inhibitor cocktail, and 1x with IDTET (0.2% Tween-20, 10 mM Tris pH 8, 0.1 mM EDTA). Samples were finally resuspended in 25 μL TT buffer (10 mM Tris pH 8 + 0.05% Tween 20) prior to on-bead library preparation. *For MITF, TFEB, and TFE3 ChIP-seq*, one million nuclei were thawed on ice and resuspended in ice-cold RLNR1 buffer (20mM Tris HCl pH 7.5, 150 mM NaCl, 1 mM EDTA, 0.5 mM EGTA, 0.4% sodium deoxycholate, 1% NP-40, 0,1% SDS, 0.5 mM DTT) + 1x protease inhibitor cocktail/PMSF. Nuclei / chromatin was sheared and recovered with the same method employed for H3K27ac. 10 μL of Dynabead Protein A and 10 μL of Protein G beads per sample were then coupled to either 2 μg of MITF antibody (ActiveMotif, 91201), 2 μg each of two TFEB antibodies (Abcam ab2636, Cell Signaling CST 37785 (D2O7D), or 2 μg each of two TFE3 antibodies (Abcam ab93808, Millipore Sigma HPA023881, or NBP1-89976). Bead/antibody complexes were added to each sample and rotated overnight at 4°C. The samples were washed with the following buffers: 2x RLNR1 + PIC/PMSF/DTT buffer, 3x LWB-RCNR1 buffer (10 mM Tris HCl pH 7.5, 1 mM EDTA, 0.7% sodium deoxycholate, 1% NP-40, 250 mM LiCl) + PIC/PMSF, 2x TET buffer, 1x IDTET buffer, and then resuspended in 25 μL TT for on-bead library preparation.

Sequencing libraries for ChIP and input samples were prepared with NEBNext Ultra II DNA library prep kit (NEB, E7645) reagents according to the manufacturer’s protocol on beads suspended in 25 μL TT (10 mM Tris/HCl pH 7.5, 0.05% Tween-20), with reagent volumes reduced by half. DNA was eluted and crosslinks reversed by adding 4 μL 10% SDS, 4.5 μL 5 M NaCl, 3 μL EDTA, 4 μL EGTA, 1 μL proteinase K (20 mg/ml), 16 μL water, incubating for 1 h at 55°C, then 30 minutes to overnight at 65°C. DNA was purified using 2 μL of SpeedBeads (Cytiva GE45152105050250) in a 20% PEG8000 / 1.5M NaCl solution to final concentration of 12% PEG, then eluted with 25 μL TT. DNA contained in the eluate was then amplified for 12-14 cycles in 25 μL PCR reactions using NEBNext High-Fidelity Q5 2X PCR Master Mix (NEB) and 0.5 mM each of primers Solexa 1GA and Solexa 1GB. Resulting libraries were size selected by gel excision to 200-500 bp, purified, and single end sequenced using an Illumina NovaSeq instrument.

### IDR analysis of ATAC and ChIP peaks

ATAC- and ChIP-seq experiments were performed in replicates with corresponding input experiments. Peaks were called using HOMER findPeaks for each tag directory with relaxed peak finding parameters ‘-L 0 -C 0 -fdr 0.9’. ATAC peaks were called with additional parameters ‘-minDist 200 -size 200’. Irreproducible Discovery Rate (IDR) (v2.0.4) was used to test for reproducibility between replicates^76^; using only peaks with an IDR <0.05 for downstream analyses. For groups with more than two replicates, peak sets from all pairwise IDR comparisons were merged into a final set of peaks for further analysis. Encode blacklist regions were removed from final peak sets using the HOMER mergePeaks command with the ‘-prefix’ parameter.

### ATAC and ChIP analysis

To quantify differential regions of H3K27 acetylation between genotypes, IDR ATAC peaks from WT and KO microglia were merged with HOMER’s mergePeaks command. Merged ATAC-seq peaks were then annotated with raw H3K27ac tag counts from each sample with HOMER’s annotatePeaks using parameters -noadj and -size 800. Subsequently, DESeq2 was used to identify the differential H3K27ac signal with the cutoffs of FC>2 and p.adj<0.05 using getDiffExpression.pl. To identify motifs enriched in differential peak regions over background, HOMER’s motif analysis (findMotifsGenome.pl) including known default motifs and de novo motifs was used. Background sequences were comprised of random GC-matched genomic sequences. For TFE3-bound promoter and enhancer analyses, promoters were defined as ±500 bp from the TSS and enhancers as outside this region, both having a minimum of ≥16 normalized merged tags for H3K27ac and ≥64 normalized tags for TFE3 IDR peaks.

### Motif co-occurrence analysis

The Motif Co-occurrence Analysis Tool, MCAT was used to discover and differentially analyse motif co-occurrence patterns between two genomic peak sets. For the analysis presented in Figure 5e, *de novo* enriched motifs were identified for the 1272 TFE3 peaks exhibiting a >2 fold increase in H3K27ac in *Sgsh* KO microglia and the 3771 TFE3 peaks exhibiting significant (>16 normalized tags) but unchanged (<1.2 FC) H3K27ac using HOMER’s findMotifsGenome.pl function^77^. Subsets of 5-8 of the most significantly enriched de novo motifs from each peak set were curated to form the target motif set. MCAT then annotated each peak with motif presence/absence over a range of motif score thresholds and generated motif co-occurrence tables for all possible unique motif combinations at each threshold (2^n^, where n is the number of motifs in target set) and compared them for significant differences. MCAT provided normalized counts for each motif combination in both peak sets and assessed statistical significance using p-values derived via standard permutation testing^123^. Different subsets of the *de novo* motifs identified in TFE3 peaks with H3K27ac^induced^ and H3K27ac^unchanged^ were iteratively evaluated to estimate contributions of individual motifs to differential representation of specific combinations in the two target peak sets. As expected, the frequencies of specific motif co-occurrences increased as motif thresholds were reduced. However, most of the frequently observed differential motif combinations were consistently identified at similar levels of significance over a broad range of motif score thresholds. The analysis presented in Fig. 5e is based on using the highest motif score threshold at which all peaks in both peak sets contain at least one of the five motifs (i.e., the null set =0).

### Antisense Oligonucleotide (ASO) treatments

ASOs were diluted in PBS without Ca2^+^ and Mg2^+^ and administered via intracerebroventricular (ICV) bolus injection as previously described. Briefly, ASOs were delivered in a 10 μL volume administered over 1 minute using the following coordinates: anteroposterior, +0.3 mm; mediolateral, +1.0 mm; and dorsoventral, -3.0 mm relative to bregma (right hemisphere). Single ASO treatment groups were administered with a 700 μg dose and combination-ASO treatment groups were administered with a mixed solution of 500 μg *Mitf*-ASO and 500 μg *Tfe3*-ASO. All ASOs used were 5-10-5 MOE gapmers (designed and provided by Ionis Pharmaceuticals, Inc.). ASOs were designed to target the intron, exon or 3’-UTR of the mouse gene of interest. ASO sequences were as follows: Control ASO: CCTATAGGACTATCCAGGAA; mouse-specific TFE3-targeting ASO: ACACTTATTTTTTTTCTTCC; mouse-specific MITF-targeting ASO: TCTTCTTTTTTTTCTCTTGA; and mouse-specific TFEB-targeting ASO: CCTTCATTTTTACCTTTACT.

### snRNA-seq analysis of human microglia

We obtained the expression data of two large publicly available datasets that performed snRNA-seq on the frontal cortex of human cohorts. The first dataset is from V Gazestani *et al*. (2023)^97^, which performed snRNA-seq on 52 surgical biopsies from patients undergoing ventriculoperitoneal shunt placement for the treatment of suspected normal pressure hydrocephalus (NPH). The second dataset is from Green *et al*. (2024)^98^, which studied single-nucleus RNA-sequencing profiles sampled from 437 older individual donors from the ROSMAP cohort. The UMAP representation and cell-type assignments were computed and extracted identically to the published results in the scanpy^124^ framework. Aggregated enrichment scores of the bulk *ex vivo* microglia signature for the *Sgsh* KO genes (8 months; fold change>2, p.adj<1e-10) for microglial states were generated using the “score_genes” function in scanpy and normalized within each dataset.

### Statistical analyses

Statistical analyses of snRNA-seq, scRNA-seq, bulk RNA-seq, and ChIP-seq data were performed as described above. All other data were analysed using Prism 10 (GraphPad). Information regarding individual statistical tests is provided in each panel’s corresponding figure legend.

## Data Availability

All raw and processed bulk sorted RNA-seq, sn/scRNA-seq, ATAC-, and ChIP-seq data are available via the NCBI GEO repository under accession numbers GSE278447, GSE278448, GSE278449, and GSE278451.

## Code Availability

All code used to analyse data in this manuscript is available upon request.

## ACKNOWLEDGEMENTS AND FUNDING

The authors would like to thank Cody Fine and Mateo Espinoza with the UCSD Human Embryonic Stem Cell Core for their equipment access and assistance with sorting. We thank the Gordts lab and Patrick Secrest for housing, maintenance, and supplying of *Sgsh* mice. We thank Amanda Roberts and Chelsea Cates with the Scripps Research Institute Animal Models Core for their assistance and *Sgsh* KO ASO injection experiments. We thank the Glass lab’s Jana Collier for requisition of all lab materials needed for this study, Katelyn Griffin for assistance with manuscript figures, and all current and previous members for their inputs. We thank Dr. Gil di Paolo and Dr. Leon Tejwani at Denali Therapeutics, and Dr. Matt Blurton-Jones and Dr. Johnny Hasselmann at UC Irvine for their valuable insights, manuscript review, and collaboration. We thank professors Chris Benner and Sven Heinz in UCSD CMM for their experimental and analysis advice on RNA and epigenetic experiments. We thank the University of California, San Diego-Cellular and Molecular Medicine Electron Microscopy Core (UCSD-CMM-EM Core, RRID:SCR_022039) for equipment access and technical assistance. The UCSD-CMM-EM Core is partly supported by the National Institutes of Health Award number S10OD023527. This manuscript includes data generated at the Institute Genomic Medicine at University of San Diego utilizing an Illumina Nova-Seq 6000 that was purchased with funding from a NIH SIG grant (#S10OD026929). We also wish to thank the UCSD Microscopy core for equipment access and technical assistance. The UCSD Microscopy core is supported by the following grant: NINDS P30NS047101. C.D.B. was supported by the National Science Foundation Graduate Research Fellowship (NSF-GRFP) under Grant no. DGE-2038238, and by the Ruth Kirschstein Institutional National Research Service Award (T32 GM008666) from the National Institute of General Medical Sciences. Any opinions, findings, and conclusions or recommendations expressed in this material are those of the author(s) and do not necessarily reflect the views of the National Science Foundation. CKG was supported by NIH grants NS096170 and AG083977, and by grants from the Cure Alzheimer Fund, the Alzheimer’s Association, and the JPB foundation. JCMS was supported by the Sanfilippo Children’s Foundation, the Fondation Sanfilippo Suisse, and the California Institute for Regenerative Medicine (CIRM) grant DISC2-13077.

## AUTHOR CONTRIBUTIONS

C.D.B, J.C.M.S, and C.K.G. conceived the project and wrote the manuscript with input from co-authors. Experiments and analysis were performed by C.D.B., J.C.M.S., A.L., L.W., N.J.S., assisted by M.P. and S.O.. Maintenance and supplying of *Sgsh* KO mice was performed in the lab of P.G.. Further experimental guidance and data analysis assistance was provided by N.J.S.. ASO development, testing/validation, and *in vivo* injections were performed by C.H., J.D., and F.K.. Motif co-occurrence analysis was performed by P.S. and C.K.G.. Immunofluorescent imaging and analysis was performed by B.L. and Y.Z.. Human snRNA-seq analyses of *Sgsh* KO signature overlap was performed by V.S. and B.S.. The project was supervised by C.K.G.. All authors contributed to editing and reviewing the manuscript.

## DECLARATION OF INTERESTS

C.K.G. is a cofounder and member of the scientific advisory board of Asteroid Therapeutics. C.H., J.D., and F.K. are employees and/or shareholders of Ionis Pharmaceuticals, Inc.

**Correspondence and requests for materials** should be addressed to and will be fulfilled by Christopher K. Glass.

## EXTENDED DATA FIGURE LEGENDS

**Extended Data Figure 1.**
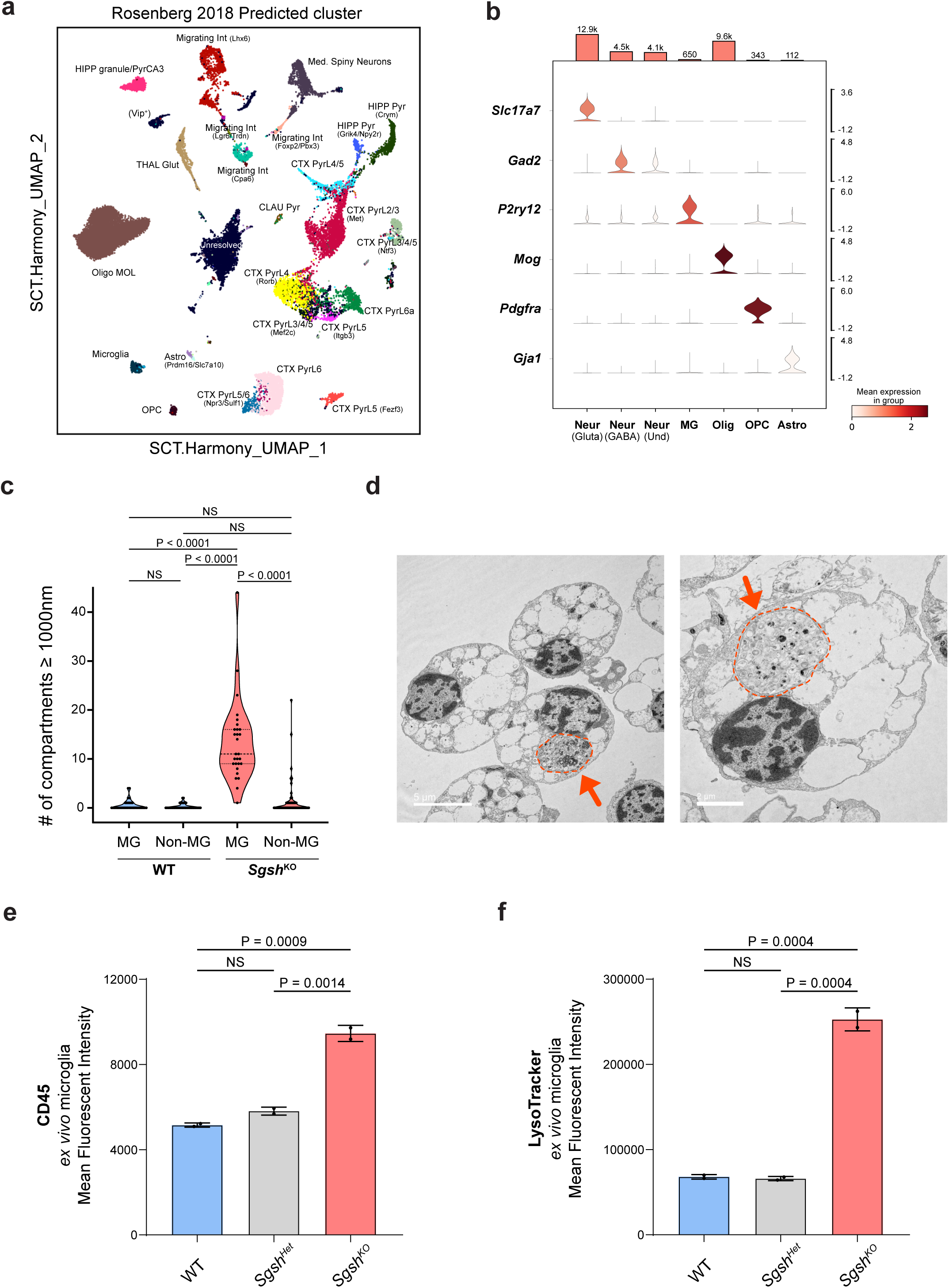
Microglia are the most severely affected cell type in brains of *Sgsh* KO mice. **(a)** UMAP plot of cell type / cluster identities after reference-based cell type label transfer and embedding projection using the reference dataset from Rosenberg *et al*.^61^. **(b)** Violin plots of cell type-specific marker genes associated with clusters illustrated in Fig. 1a. **(c)** Violin plot illustrating the numbers of cytoplasmic compartments >1000 nm in microglia and non-microglial (neurons, oligodendrocytes, astrocytes) cells in WT and *Sgsh*KO mice (*n=3* biological replicates/genotype). **(d)** EM images of sorted microglia containing intracellular inclusions resembling dystrophic neurites. **(e)** Bar plots of CD45 mean fluorescence intensity values following flow cytometry in WT, Het, and *Sgsh* KO microglia derived from 8-month-old mice (*n=2* biological replicates/genotype). **(f)** Bar plots of Lysotracker Red mean fluorescence intensity values following flow cytometry in WT, Het, and *Sgsh* KO microglia from 8-month-old mice (*n=2* biological replicates/genotype). For bar plots, data are represented as mean ± standard deviation, with p values determined by one-way ANOVA followed by post-hoc Tukey for multiple comparisons.

**Extended Data Figure 2.**
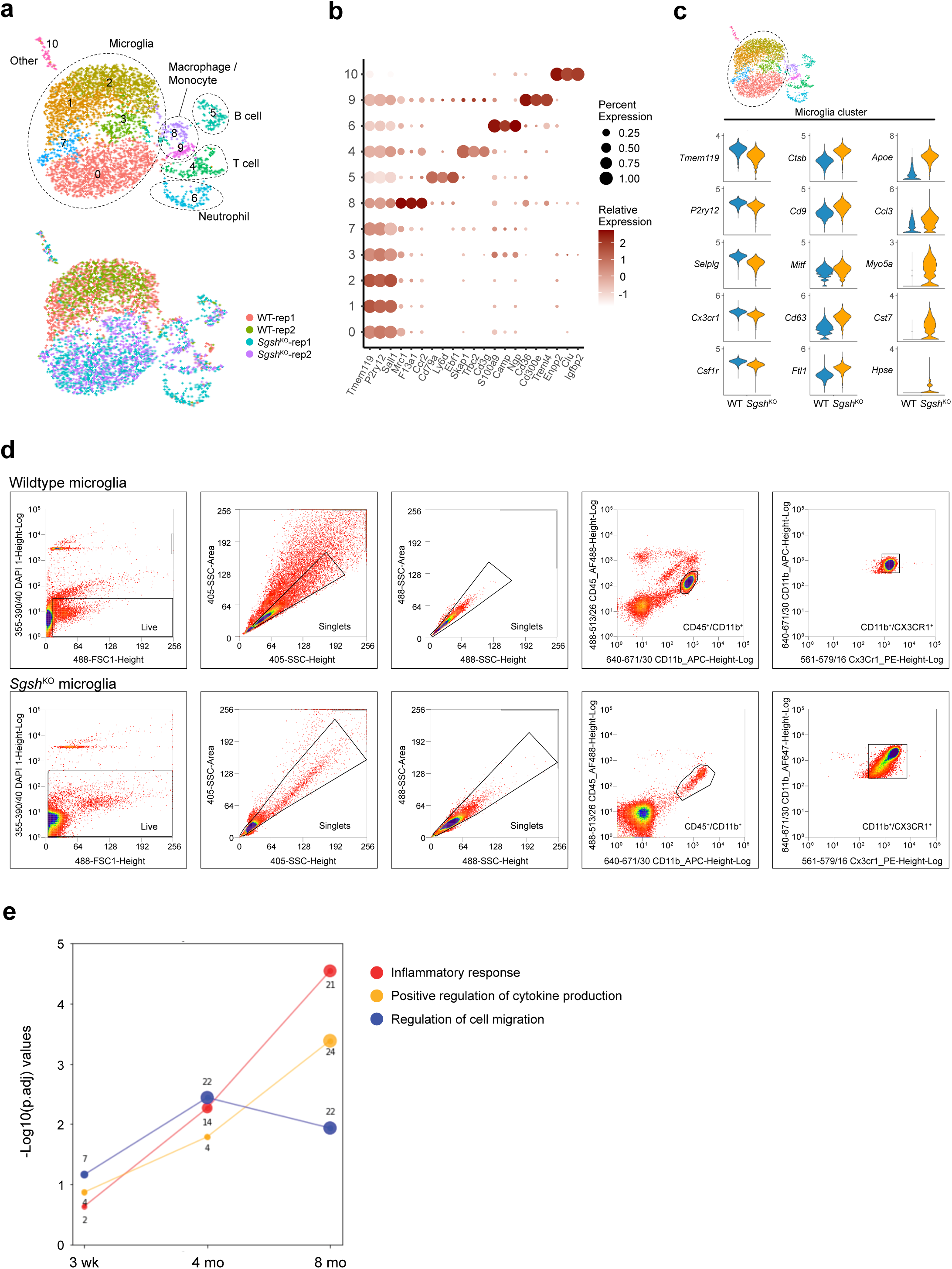
Gene expression signature of *Sgsh* KO microglia. **(a)** UMAP of clusters identified among all high-quality CD45^+^ cells of the scRNA-seq dataset (top) and individual biological replicate samples (bottom). **(b)** Dot plot of marker genes for different clusters. **(c)** Volcano plot illustrating DEGs derived from differential expression analysis of WT vs *Sgsh* KO microglia in the microglia subcluster. **(d)** Gating strategy for sorting live microglia for downstream assays. **(e)** Gene count and enrichment values for the indicated terms derived from DEGs in WT vs *Sgsh* KO microglia isolated from mice at 3 weeks, 4 months and 8 months of age.

**Extended Data Figure 3.**
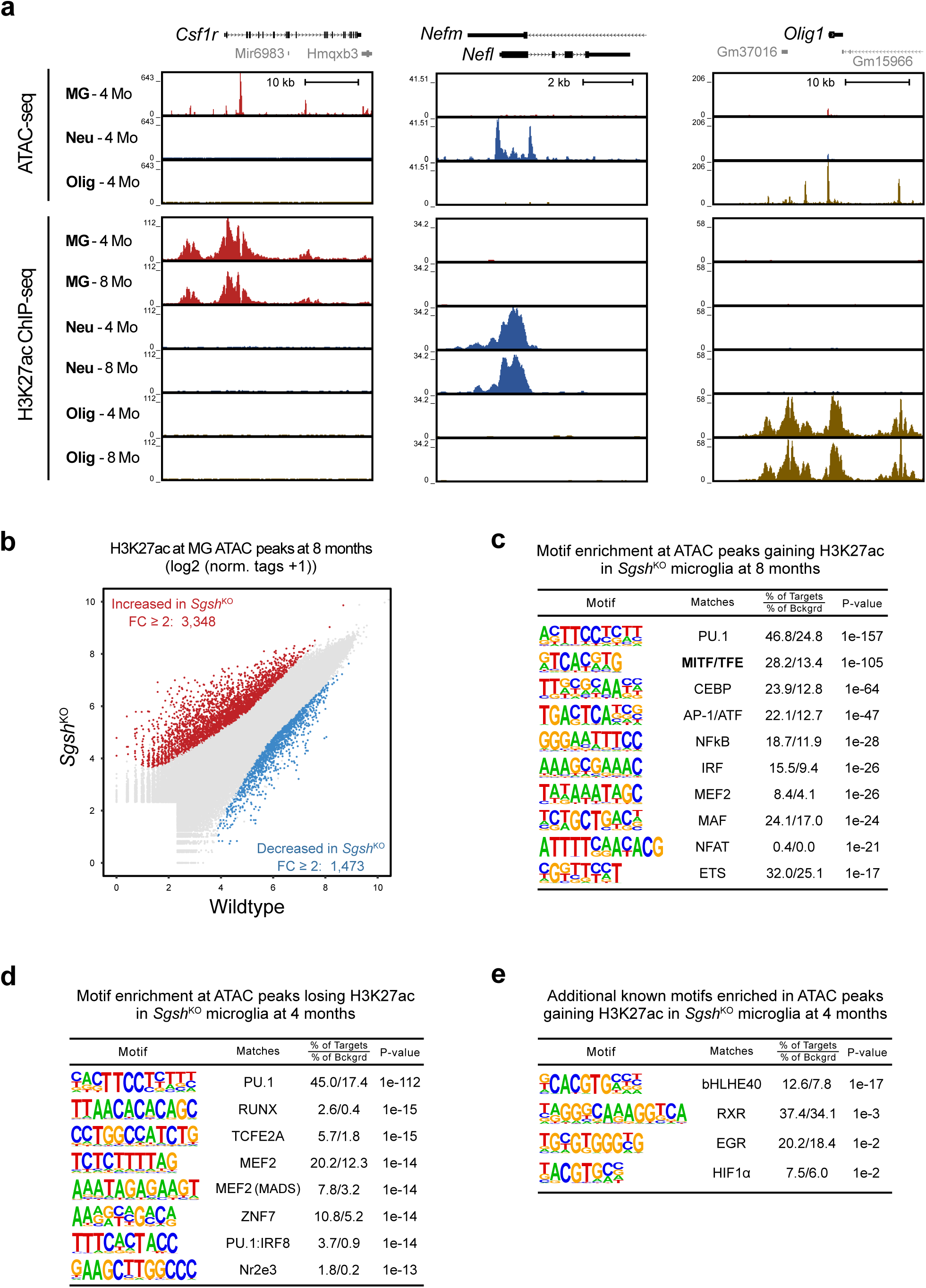
**Cell-specific effects of *Sgsh* KO on the epigenetic landscapes of microglia, neurons, and oligodendrocytes**. **(a)** Representative UCSC Genome Browser tracks for ATAC-seq and H3K27ac ChIP-seq data at cell type-specific gene loci documenting purity of each cell population (*n=2* biological replicates per genotype and per timepoint in each cell type). **(b)** Scatter plot of mean H3K27ac (log_2_(norm.tags+1)) at ATAC-seq peaks in WT and *Sgsh* KO microglia isolated at 8 months of age. Red: increased FC>2 (p.adj<0.05) in *Sgsh* KO microglia, blue: decreased FC>2 (p.adj<0.05) in *Sgsh* KO microglia. **(c)** *De novo* enrichment analysis of ATAC peaks exhibiting >2-fold increase in H3K27ac (p.adj<0.05) in *Sgsh* KO microglia at 8 months of age. **(d)** Motif enrichment at ATAC-peaks exhibiting reduced H3K27ac (>2-fold, p.adj<0.05) in *Sgsh* KO microglia compared to WT microglia at 4 months of age. **(e)** Known motifs enriched at ATAC peaks gaining H3K27ac in *Sgsh* KO microglia compared to WT microglia at 4 months of age.

**Extended Data Figure 4.**
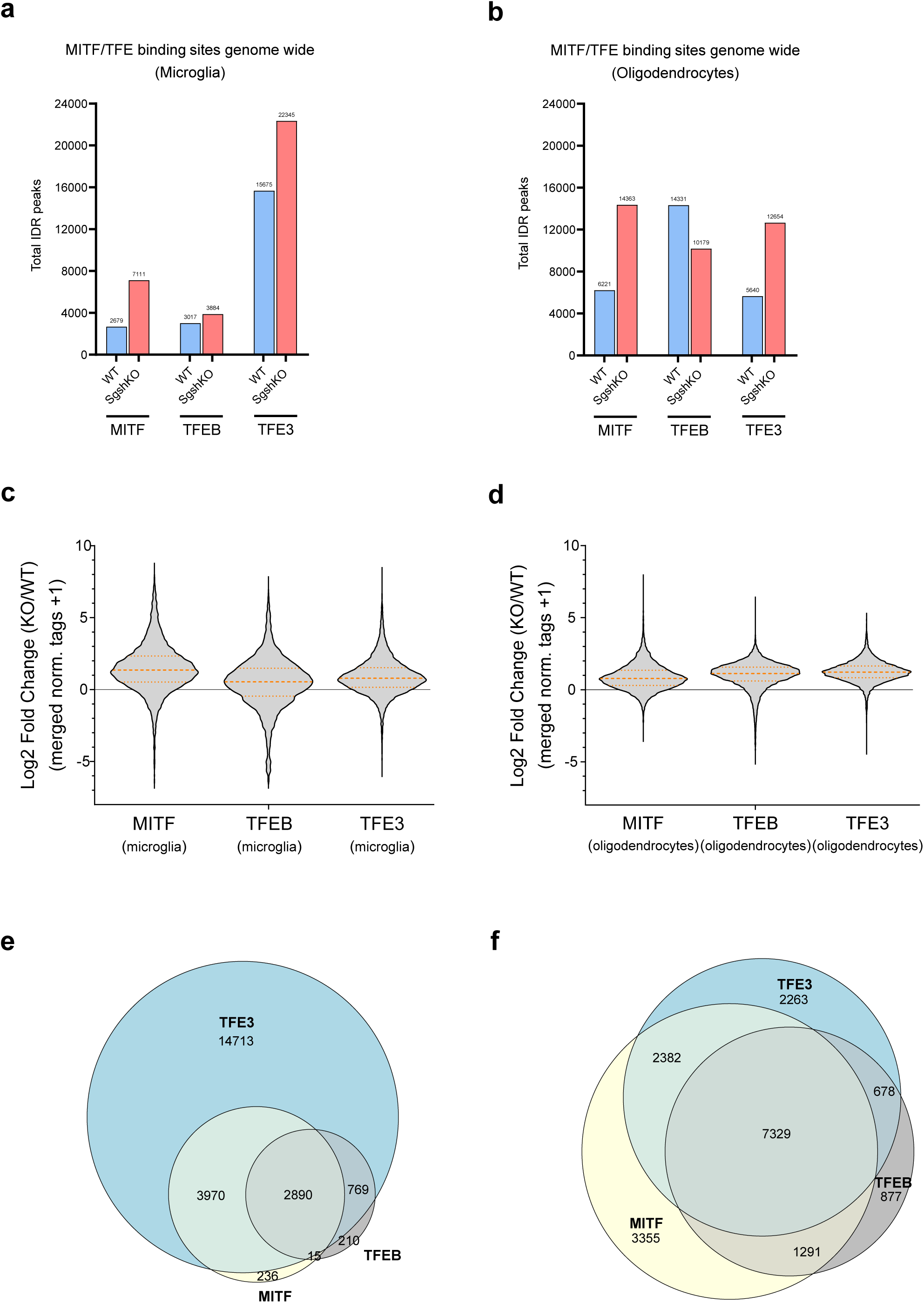
Cell-specific and genotype-dependent binding profiles of MITF, TFEB and TFE3 in WT and *Sgsh* KO microglia and oligodendrocytes. **(a)** Bar graphs of ChIP-seq peaks for MITF, TFEB, and TFE3 in microglia from WT and *Sgsh* KO mice as determined by IDR analysis of replicate samples. **(b)** Bar graphs of ChIP-seq peaks for MITF, TFEB, and TFE3 in oligodendrocytes from WT and *Sgsh* KO mice as determined by IDR analysis of replicate samples. **(c)** Violin plots of the Log_2_ fold change of ChIP-seq normalized tag counts for the indicated factors in microglia comparing *Sgsh* KO to WT mice. **(d)** Violin plots of the Log_2_ fold change of ChIP-seq normalized tag counts for the indicated factors in oligodendrocytes comparing *Sgsh* KO to WT mice. **(e)** Venn diagram indicating overlap of MITF, TFEB and TFE3 peaks in *Sgsh* KO microglia. **(f)** Venn diagram indicating overlap of MITF, TFEB and TFE3 peaks in *Sgsh* KO oligodendrocytes.

**Extended Data Figure 5.**
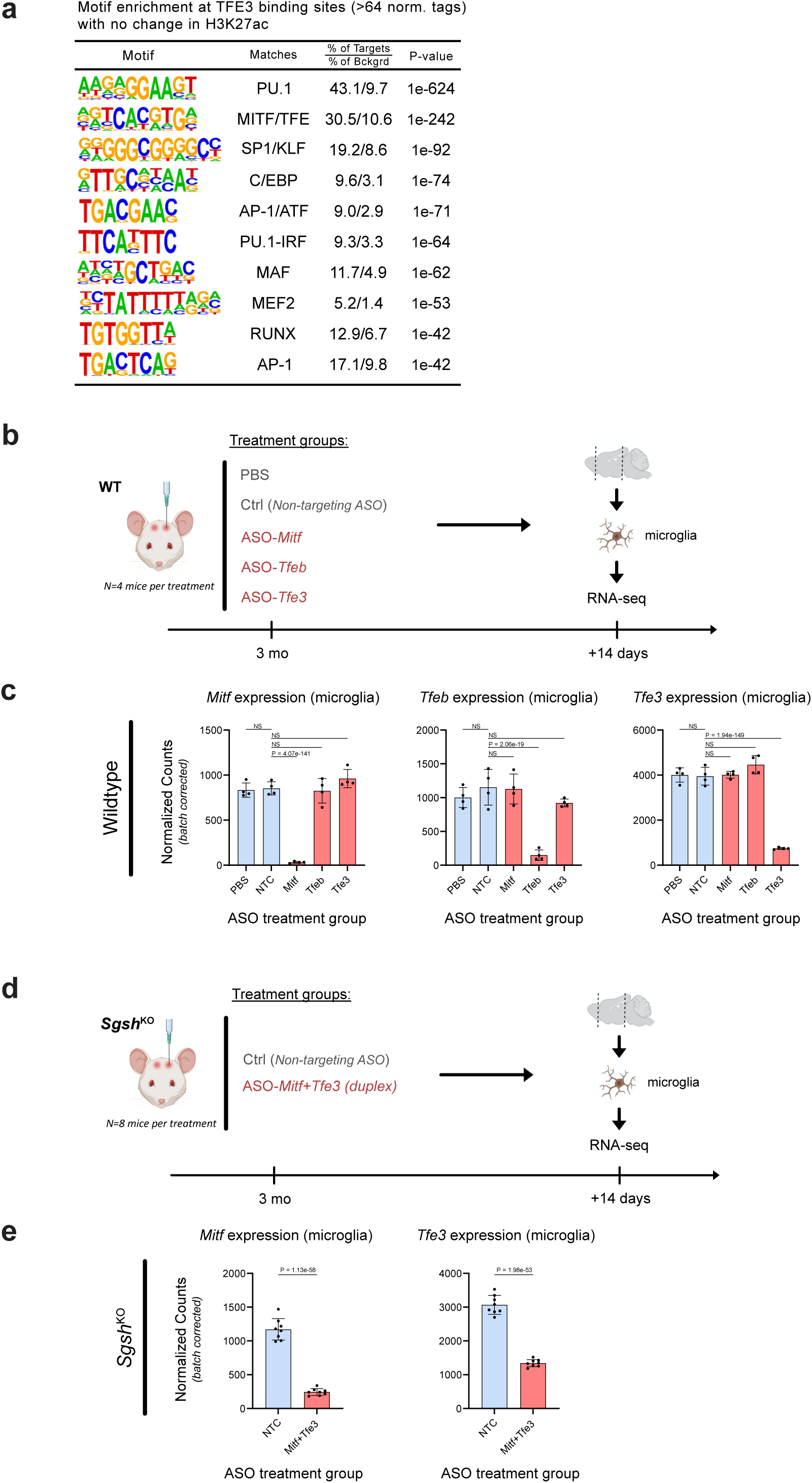
**Homeostatic and *Sgsh* KO-dependent binding programs of MITF, TFEB and TFE3 in microglia and molecular functions of MITF, TFEB and TFE3 in *Sgsh* KO microglia**. **(a)** *De novo* motif enrichment of TFE3 peaks exhibiting >64 tags at locations exhibiting more than 16 normalized tags of H3K27ac in WT MG and less than ±1.2-fold change in *Sgsh* KO microglia. **(b)** Experimental design for injection of ASOs and recovery of microglia for RNA-seq analysis in WT mice. **(c)** Knockdown efficiencies for each ASO target in wildtype mice (*n=4* biological replicates/ treatment group). **(d)** Experimental design for injection of ASOs and recovery of microglia for RNA-seq analysis in *Sgsh* KO mice. **(e)** Knockdown efficiencies for each ASO target in *Sgsh* KO mice (*n=8* biological replicates/ treatment group).

**Extended Data figure 6.**
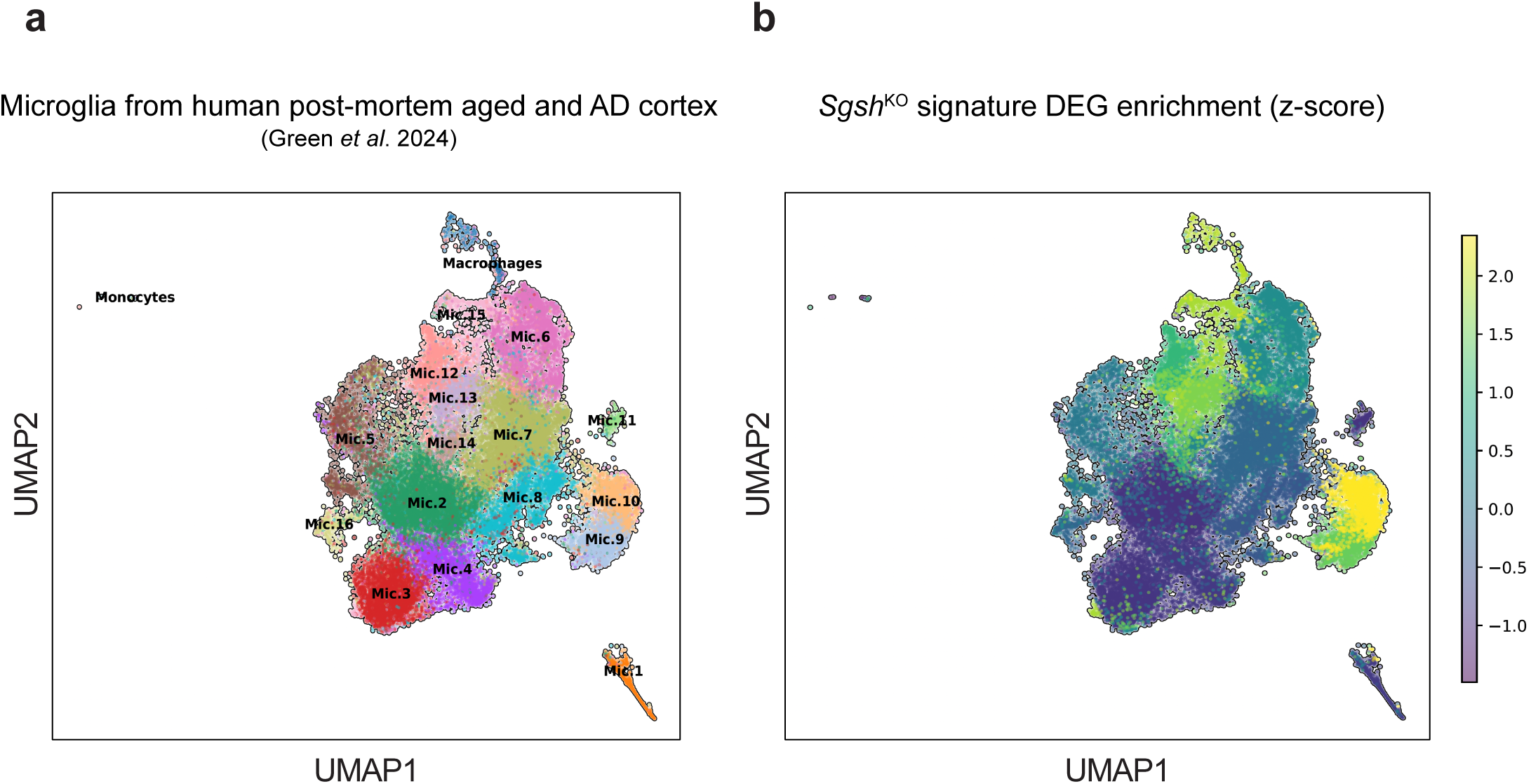
Relationships to models of age-related neurodegeneration and human microglia in AD. **(a)** UMAP plot of the cell clusters of brain myeloid cell clusters obtained from snRNA-seq of human post-mortem tissue samples from individuals with varying degrees of amyloid and tau pathology as published in Green *et al*., 2024. Each dot represents a single cell. **(b)** UMAP plot of the composite averaged z-score from human orthologs of a representative *Sgsh* KO signature gene set across all cells in the UMAP shown in **a**.

## SUPPLEMENTARY TABLE LEGENDS

**Supplementary Table 1. Bulk RNA-seq of sorted *ex vivo Sgsh* KO microglia.** DEG statistics between WT and *Sgsh* KO *ex vivo* microglia at 3 weeks, 4 months, and 8 months of age.

**Supplementary Table 2. ATAC, H3K27ac and MITF/TFE ChIP-seq of sorted *ex vivo Sgsh* KO microglia.** Merged (WT and Sgsh KO) IDR ATAC peak regions annotated with normalized merged tag directories from ATAC-seq, H3K27ac ChIP-seq, and MITF, TFEB, and TFE3 ChIP-seq of WT and *Sgsh* KO microglia.

**Supplementary Table 3. Sequencing and QC metrics of bulk RNA-seq experiments** A summary of sequencing and QC metrics for bulk RNA-seq experiments of sorted *ex vivo* microglia experiments.

**Supplementary Table 4. Sequencing and QC metrics of ATAC- and ChIP-seq experiments** A summary of sequencing and QC metrics for ATAC-, H3K27ac, and transcription factor ChIP-seq experiments of microglia, neurons, and oligodendrocytes.

